# Genome and time-of-day transcriptome of *Wolffia australiana* link morphological extreme minimization with un-gated plant growth

**DOI:** 10.1101/2020.03.31.018291

**Authors:** Todd P. Michael, Evan Ernst, Nolan Hartwick, Philomena Chu, Douglas Bryant, Sarah Gilbert, Stefan Ortleb, Erin L. Baggs, K. Sowjanya Sree, Klaus J. Appenroth, Joerg Fuchs, Florian Jupe, Justin P. Sandoval, Ksenia V. Krasileva, Ljudmylla Borisjuk, Todd C. Mockler, Joseph R. Ecker, Robert A. Martienssen, Eric Lam

## Abstract

Wolffia is the fastest growing plant genus on Earth with a recorded doubling time of less than a day. Wolffia has a dramatically reduced body plan, primarily growing through a continuous, budding-type asexual reproduction with no obvious phase transition. Most plants are bound by the 24-hour light-dark cycle with the majority of processes such as gene expression partitioned or phased to a specific time-of-day (TOD). However, the role that TOD information and the circadian clock plays in facilitating the growth of a fast-growing plant is unknown. Here we generated draft reference genomes for *Wolffia australiana* (Benth.) Hartog & Plas to monitor gene expression over a two-day time course under light-dark cycles. *Wolffia australiana* has the smallest genome size in the genus at 357 Mb and has a dramatically reduced gene set at 15,312 with a specific loss of root (WOX5), vascular (CASP), circadian (TOC1), and light-signaling (NPH3) genes. Remarkably, it has also lost all but one of the NLR genes that are known to be involved in innate immunity. In addition, only 13% of its genes cycle, which is far less than in other plants, with an overrepresentation of genes associated with carbon processing and chloroplast-related functions. Despite having a focused set of cycling genes, TOD cis-elements are conserved in *W. australiana*, consistent with the overall conservation of transcriptional networks. In contrast to the model plants *Arabidopsis thaliana* and *Oryza sativa*, the reduction in cycling genes correlates with fewer pathways under TOD control in Wolffia, which could reflect a release of functional gating. Since TOD networks and the circadian clock work to gate activities to specific times of day, this minimization of regulation may enable Wolffia to grow continuously with optimal economy. Wolffia is an ideal model to study the transcriptional control of growth and the findings presented here could serve as a template for plant improvement.

## Introduction

*Wolffia* has the distinction of being the duckweed genus with the smallest (*Wolffia angusta*) as well as the fastest growing (*Wolffia microscopica*) species of known flowering plants (Sree et al. 2015b). Plants belonging to this genus are highly reduced in their morphology and anatomy, lacking roots and containing only the green floating frond, which is essentially a fused leaf and stem without any vasculature (Figure 1). Wolffia typically measure only a few millimeters to less than a millimeter in size (Landolt 1986) and grow as colonies of two individuals, one mother frond budding and giving rise to one or more daughter fronds (Figure 1A). Anatomically, however, at least four different generations of plants (total of 10-14 individuals) can be found within one colony, highlighting their adaptive preparedness for fast vegetative multiplication (Sree et al. 2015a; Bernard et al. 1990; Lemon and Posluszny 2000) (Figure 1C, Suppl Video).

**Figure 1.**
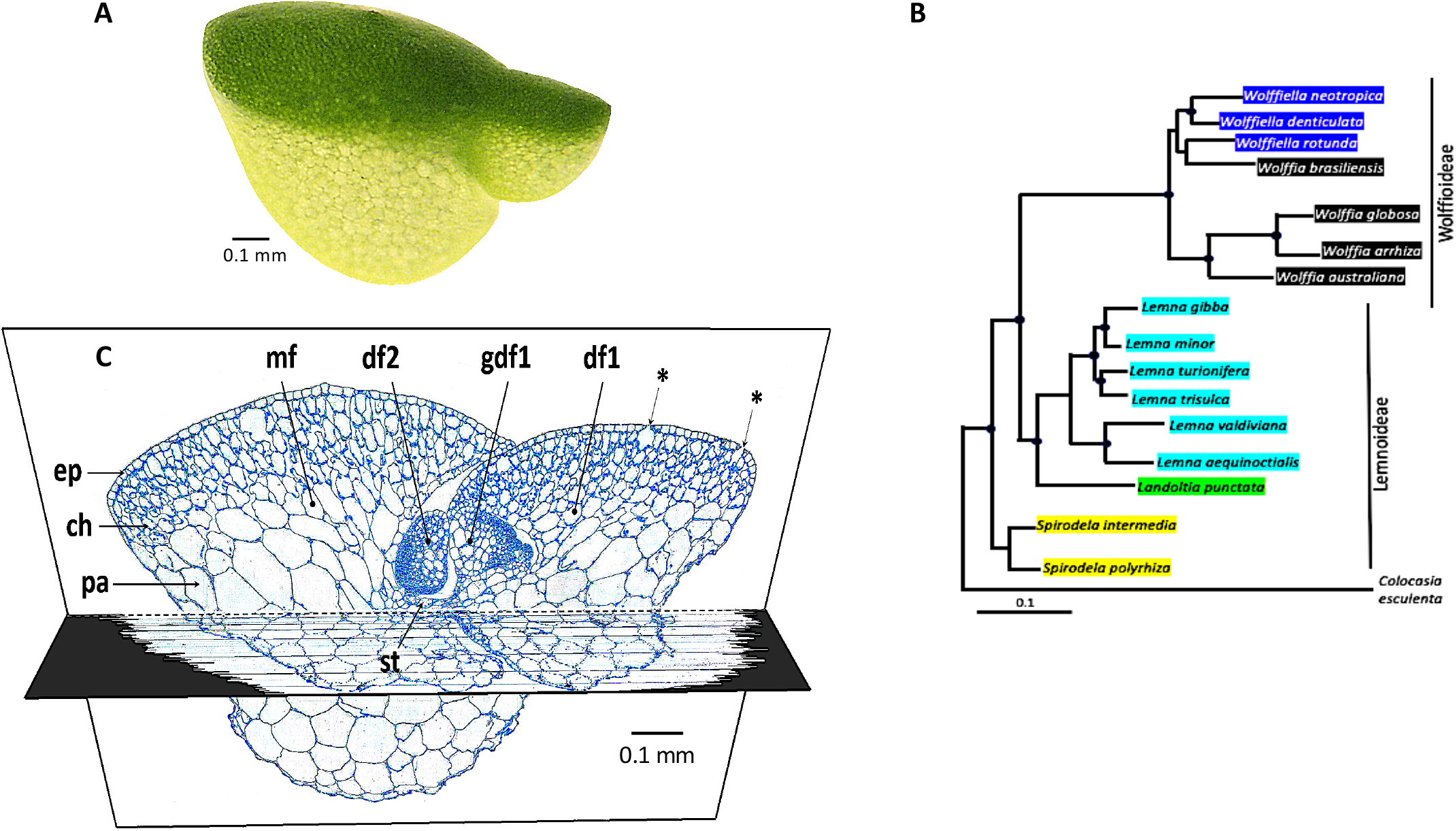
Wolffia is a simple plant with a limited number of cells and structures. **A)** Brightfield image of wa8730. A video with time tracking of the asexual propagation of wa7733 in culture can be found in the Supplemental Information. **B)** Phylogenetic relationship between representative species from the 5 genera of duckweed. Each color represents a distinct genus. Modified from (Borisjuk et al. 2015). **C)** Cross section (1 um thick section) of *W. australiana* stained with methylene blue. **mf**: mother frond; **df**: daughter frond; **gdf**: grand-daughter frond; **ep**: epidermal cells; **ch**: chlorenchyma cells; **pa**: parenchyma cells; **asterisks** indicate some of the stomata; **st**: stipe tissue.

Of applied interest is the high nutritional quality of Wolffia for human and livestock food. *Wolffia globosa* has been traditionally eaten in Southeast Asian countries like Thailand, Vietnam, Laos and Cambodia. Not only is the protein content high (~30% dry weight), the level for different amino acids is well above the recommendation of the World Health Organization (WHO) (Appenroth et al. 2018). Although the fatty acid content is low, its composition is suitable for the human diet. Interestingly, the mineral content can be modulated as per the desired content (Appenroth et al. 2018). Additionally, it was found that the plant extracts did not possess any cytotoxic or anti-proliferative effects on tested human cell lines (Sree et al. 2019). Taken together with the high growth rate, *Wolffia* has the added potential to be biofortified and developed as a more nutritious food crop for the future.

Wolffia is part of a family of aquatic non-grass monocots known as Duckweeds (Lemnaceae) (Figure 1B). There are five genera of duckweed (Spirodela, Landoltia, Lemna, Wolffiella and Wolffia), and their genome sizes span an order of magnitude (Wang et al. 2011; Hoang et al. 2019; Bog et al. 2015). The Greater Duckweed, *Spirodela polyrhiza*, which is most basal with the largest body size and most complex organization, has the smallest genome at 158 megabase (Mb), while Wolffia, the most derived genus of the Duckweed family, has genomes that span from *W. australiana* at 357 Mb to *W. arrhiza* at 1,881 Mb. *S. polyrhiza* (sp7498) was the first duckweed genome to be sequenced, which revealed a reduced set of protein coding genes at 19,623 that are conserved across flowering plants (Wang et al. 2014; An et al. 2019). A chromosome-resolved genome for a second accession sp9509 revealed that in addition to a reduced gene set, the Spirodela genome displayed low intraspecific variance and also has highly reduced ribosomal arrays and cytosine methylation, consistent with the genome being specialized for growth (Hoang et al. 2018; Michael et al. 2017).

Since Wolffia is the fastest growing angiosperm with a doubling time of as little as 18 hours (S. Sree, unpublished data), we wanted to understand what special adaptations to the genome enabled this rapid growth. Therefore, we chose *W. australiana*, since it has the smallest genome of this genus, for whole genome sequencing and time-of-day (TOD) expression profiling. Green organisms from algae to higher plants partition their biology to coincide with the light-dark cycle, which enhances their ability to anticipate changing conditions (Michael et al. 2003; Dodd et al. 2005; Green et al. 2002). In the model plant *Arabidopsis thaliana*, as much as 90% of its genes are expressed, or phased, to a specific TOD to optimize growth and this global transcriptional regulation is conserved across higher plants (Michael et al. 2008a, 2008b; Filichkin et al. 2011). Underlying TOD-specific expression is the circadian clock that modulates, or “gates”, an organism’s response to the environment, but also continues to oscillate with a period of about 24 hours (circa diem, circadian) in the absence of timing cues (McClung 2019). Specifically, different aspects of growth such as cell expansion and the cell cycle are regulated by the circadian clock in concert with environmental signals (Fung-Uceda et al. 2018).

Here we report the draft genomes for two *W. australiana* accessions wa7733 and wa8730, which have doubling times of 1.56 and 1.66 days, respectively (Table 1; Figure S1), consistent with the previous measurement of 1.39 days (Sree et al. 2015b). Compared to *S. polyrhiza*, the Wolffia draft genomes have a further reduced number of core eukaryotic protein coding genes consistent with a minimal body plan lacking elaborate structures and roots. A two-day time course for the TOD expression profile revealed a limited number of cycling genes focused almost exclusively on photosynthesis suggesting Wolffia may have unrestricted, or ungated growth. Together with its budding-like replication strategy, Wolffia is thus similar to a “green yeast,” with its reduced body plan, rapid growth, and minimized set of core eukaryotic genes.

**Table 1.** *Wolffia australiana* growth rates. Growth rates of four clones of *Wolffia australiana* grown in N-medium at 25°C and continuous white light intensity of 100 μmol m^−2^ s^−1^ for 7 days. RGR: Relative Growth Rate, DT: Doubling Time and RY: Relative (weekly) Yield.

| **Clone No.** | **RGR (d<sup>-1</sup>) ± SE** | **DT (d) ± SE** | **RY (g g<sup>-1</sup> Week<sup>-1</sup>) ± SE** |
| --- | --- | --- | --- |
| wa7211 | 0.3739±0.0068 | 1.857±0.033 | 13.78±0.68 |
| wa7540 | 0.4890±0.0062 | 1.419±0.018 | 30.81±1.37 |
| wa7733 | 0.4454±0.0118 | 1.562±0.044 | 22.96±1.75 |
| wa8730 | 0.4167±0.0485 | 1.66±0.02 | 18.54±0.63 |

RGR: relative growth rate; DT: doubling time; RY: relative yield

## Results

### Wolffia genome

*W. australiana* has the smallest reported genome in the Wolffia genus with an estimated genome size of 375 Mb for accession wa8730 and 357 Mb for accession wa7733 (Wang et al. 2011). Both accessions are from Australia with wa7733 from Mount Lofty Range, Torrens Gorge in South Australia (S 138, E 766) and wa8730 from Singleton, Doughboy Hollow, New South Wales (S 138, E 766). We sequenced and *de novo* assembled wa7733 and wa8730 using Pacific Bioscience (PacBio) single molecule real-time sequencing (SMRT). We also generated BioNano optical maps to correct contigs and help initial scaffolding of the assemblies (Table S1), which resulted in final assemblies of 393 and 354 Mb, with longest scaffolds of 5.3 and 1.7 Mb, and scaffold N50 lengths of 836 and 109 Kb for wa7733 and wa8730, respectively (Table 2). The assembled genomes sizes are consistent with that predicted by kmer (k=19) frequency analysis (Figure S2), but smaller than predicted by new flow cytometry estimates possibly reflecting missing high copy number repeat sequence (Table S2; Methods).

**Table 2.** Genome assembly statistics.

|  | **wa7733** | **wa8730** | **wa7450\*** | **sp9509** | **sp7498** |
| --- | --- | --- | --- | --- | --- |
| Genome size estimate flow cytometry (Mb) | 432 | 441 | 432 | NA | 159 |
| Genome size estimate kmer19 (Mb) | 343 | 342 | 345 |  | 185 |
| Genome final assembly (bp) | 393,842,592 | 354,558,446 | 336,441,020 | 138,592,155 | 145,274,398 |
| Contig (#) | 2,578 | 5,250 | 163,572 | 20 | 21 |
| Genome contig assembly size (bp) | 359,766,217 | 337,899,876 | NA | NA | NA |
| Longest contig (bp) | 1,664,978 | 679,034 | 128,983 | NA | NA |
| N50 contig length (bp) | 256,298 | 102,418 | 15,364 | NA | NA |
| L50 contig (#) | 404 | 1,000 | 6,346 | NA | NA |
| Longest scaffold (bp) | 5,333,369 | 1,714,878 | 188,874 | 11,560,055 | 12,728,324 |
| N50 scaffold length (bp) | 836,551 | 109,493 | 17,957 | 7,949,387 | 8,107,549 |
| L50 scaffold (#) | 122 | 753 | 5,404 | 8 | 8 |
| BUSCO complete scaffolds (%) | 69 | 70 | NA | 79 | 78 |
| Full length LTRs (#) | 2,510 | 1,892 | NA | 801 | 567 |
| Masked (%) | 50 | 50 | NA | 26 | 25 |
| Protein coding genes (#) | 15,312 | 14,324 | NA | 17,510 | 17,057 |
| cDNA mapping (%) | 100 | 100 | NA | NA | NA |
| Coverage PacBio (fold) | 91 | 45 | NA | NA | NA |
| Coverage Illumina (fold) | 80 | 68 | 62 | NA | NA |
| Mapping PacBio (%) | 98 | 96 | NA | NA | NA |
| Mapping Illumina (%) | 98 | 95 | 97 | NA | NA |

\*wa7450 assembled with spades and Illumina reads only
wa7733 and wa8730 assembled with Canu
NA, not available

We checked the completeness of these assemblies by mapping 1,780 high confidence Sanger sequenced *W. australiana* cDNA clone sequences deposited at Genbank by the Waksman Student Scholar program (WSSP). We found that 100 and 99.7% of the cDNAs mapped to the wa7733 and wa8730 genomes, respectively. Also, the PacBio reads used for the assembly and the Illumina reads used for polishing, had a >95% mapping rate to their respective assemblies, consistent with the lack of contamination (bacterial) in the sequencing data and the completeness of the assemblies (Table 2). The contiguity (contig N50 length) of the Wolffia genomes is modest in these draft assemblies, which makes chromosome scale comparisons more challenging, but more than sufficient to capture the gene space for understanding growth related and TOD-gated pathways.

The *W. australiana* has the smallest genome in the Wolffia genus (Wang et al. 2011; Hoang et al. 2019), which suggests that like Spirodela it has a reduced amount of repeat sequence, since repeat sequence is what drives genome size differences in plants (Michael and VanBuren 2020). Therefore, we looked at ribosomal DNA (rDNA) and long terminal repeats (LTRs) in the *W. australiana* genomes to understand why it is smaller than others in the genus. It has been shown that *W. australiana* has one 45S (18S, 5.8S, 26S) and two 5S rDNA loci (Hoang et al. 2019). Only a limited amount of the 45S and 5S rDNA were captured in the assemblies (Table S3), which is expected due to the similarities in these repeats. The wa7733 assembly captured double that of the wa8730 assembly consistent with the higher contiguity in the former and possibly reflecting their differences in final scaffolded assembly size (Table 2). To estimate the number of rDNA repeats in each genome we performed a coverage analysis with the Illumina reads and found that they had a reduced repeat content with ~200 copies, double that of Spirodela but still half of the number found in Arabidopsis (Table S3). Next, we identified full-length, intact LTRs across both genomes and found 2,510 and 1,892 for wa7733 and wa8730 respectively, which was three times more than Spirodela (Table 2). Repeats (solo + intact) made up over 50% of both of the Wolffia genomes, which is twice the repeat content found in Spirodela (Michael et al. 2017). While Spirodela has a high solo::intact ratio (8.2) (Michael et al. 2017), Wolffia has an even higher ratio at 11 to 14 in wa7733 and wa8730 respectively, consistent with *W. australiana* more actively purging its TEs leading to a much smaller sized genome than other Wolffia species (Wang et al. 2011).

The two draft Wolffia genomes are highly collinear with one another at the gene level but there are examples of structural changes and gene loss and gain (Figure 2A, B; Figure S3). Most of the structural variations are associated with insertion/deletions (INDELs) between 50 and 500 bp (Figure S4). In addition, both Wolffia genomes are largely collinear with Spirodela (Figure 2; Figure S5). One specific example is in the evolutionarily conserved linkage between core circadian clock genes *LATE ELONGATED HYPOCOTYL* (*LHY*) and *PSEUDO RESPONSE REGULATOR 37* (*PRR37*) on Chr20 of the sp9509 genome that is found as far back as bryophytes and shared across other dicots and non-grass monocots (Figure 2B). While the genic regions of the Wolffia genomes are mostly collinear with those of the Spirodela genomes, the intergenic and repeat sequences are expanded in the Wolffia genomes explaining its increased genome sizes (Figure 2; Figure S3). While the *W. australiana* genomes are larger than the Spirodela genome due to increased repeat content, like the Spirodela genome they are likely smaller than the other Wolffia genomes in the genus due to active purging of LTRs as evidenced by the high solo::intact ratio.

**Figure 2.**
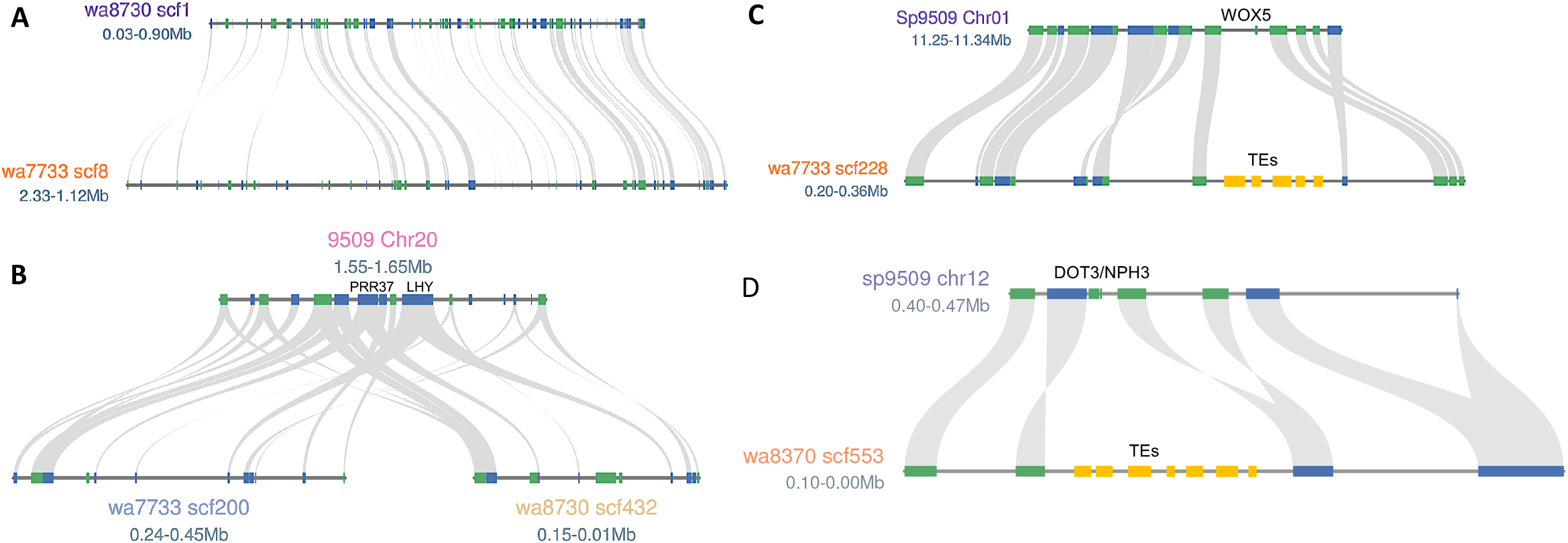
The Wolffia genome is collinear with Spirodela with bloating and loss of genes due to TEs. **A)** wa8730 scf1 is collinear with wa7733 scf8. **B)** Conserved core circadian clock genes PRR37 and LHY on Spirodela chromosome 20 (Chr20) are collinear in wa8730 scf200 and wa7733 scf432. **C)** WOX5 is lost in Wolffia compared to Spirodela due to TE insertions (yellow boxes). **D)** DOT3/NPH3 loci in Spirodela lost in Wolffia due to LTR insertions. Grey lines represent syntenic connections. Green and blue boxes are forward and reverse representation for genes, respectively. Yellow boxes are transposable elements (TEs).

### Comparative BUSCO analysis reveals genes missing in Wolffia

As an additional check of genome assembly completeness we leveraged BUSCO (**<u>B</u>**enchmarking **<u>U</u>**niversal **<u>S</u>**ingle-**<u>C</u>**opy **<u>O</u>**rthologs) (Simão et al. 2015), which searches the genome for near-universal single-copy orthologs to establish how much of the gene space has been properly assembled. We found that wa7733 and wa8730 contained 69% and 70% complete BUSCOs respectively (Table 2; Table S4). The BUSCO scores for percent complete were lower than we had expected based on the contiguity (N50 length and longest contigs) as well as the cDNA and read mapping results. Therefore, we compared the missing BUSCO genes between the two Wolffia assemblies and the two previously published high quality Spirodela genomes sp9509 and sp7498 (Wang et al. 2014; Michael et al. 2017; Hoang et al. 2018), and found that these "missing genes" were not independent across the four duckweed assemblies (Figure S6). A BUSCO gene found to be missing in one of the duckweeds is likely to be found missing in the other duckweeds. These results suggest that at least among these sequenced duckweed genomes some subset of the near-universal single-copy orthologs found in land plants do not exist in this aquatic plant family, which is consistent with the lower gene counts in duckweed (Wang et al. 2014; Michael et al. 2017; Hoang et al. 2018). Furthermore, there exists some subset of the near-universal single-copy orthologs found in land plants and Spirodela that do not exist in Wolffia.

There is a formal possibility that some of the missing BUSCO genes represent important genes for land plants that are lost in the Wolffia genome. Of the 762 and 731 missing BUSCO genes in wa7733 and wa8730 respectively, 574 (76-79%) genes were shared between them (Figure S6; Table S5). Two groups of interesting genes emerged from this list of 574 genes; first, genes that are involved with root initiation and development, and second, genes associated with cell fate and gravitropism. Of particular interest was the loss of WUSCHEL related homeobox 5 (WOX5), which is a homeodomain transcription factor responsible for root stem cell maintenance in the meristem (Sarkar et al. 2007). Since Spirodela has roots, we indeed found a likely ortholog of WOX5, while the Wolffia version of this gene has been lost and in its expected genomic location are LTRs suggesting a mechanism by which this gene may have been lost (Figure 2C). Moreover, Wolffia is also missing TOPLESS-related 2 (TPL2), which through chromatin-mediated repression specifies where stem cell daughters will exit stem cell fate in Arabidopsis (Pi et al. 2015). In addition, Wolffia is missing genes associated with stem cell fate (BLISTER, FEZ) (Schatlowski et al. 2010; Willemsen et al. 2008), mediator complex (MED3, MED9, MED33) (Mathur et al. 2011), and gravitropism (LAZY1) (Yoshihara and Spalding 2017), which together are consistent with modified signaling and transcriptional cascades for a rootless, organless plant. Since Wolffia does not contain roots, the lack of genes specifying root growth and development provides a compelling reason why these BUSCO genes are missing.

### Wolffia has a reduced set of core plant genes

We predicted genes and performed gene family analysis on the two Wolffia genomes and the two published Spirodela genomes using a standardized pipeline to ensure consistency. Similar to Spirodela, we found that Wolffia also had reduced gene sets in spite of their larger genome sizes. The wa7733 and wa8730 genome assemblies contain 15,312 and 14,324 predicted protein-coding genes respectively (Table 2), which is several thousand genes less than that found in the Spirodela genome, and this result suggests that a reduced gene set could be a common feature of duckweeds (Wang et al. 2014; Michael et al. 2017). To understand the gene families within the duckweed genomes we compared the four duckweed gene sets to 28 proteomes from complete genomes spanning algae, non-seed plants, monocots and dicots (PLAZA v4 monocots) (Van Bel et al. 2018). A multidimensional scaling (MDS) plot of the orthogroups (OGs) placed Wolffia and Spirodela next to the non-grass monocots and close to the grasses consistent with their evolutionary position (Figure S7) (One Thousand Plant Transcriptomes Initiative 2019). Interestingly, they were nested between dicot crops, basal plants and a tree, and distant from non-seed plants and algae, consistent with having a core set of higher plant genes (Wang et al. 2014).

Almost all of the Spirodela and Wolffia genes were found in OGs (93-98%; Table S6), with the majority (33-41%) having only one gene per OG (Figure S8), which means that duckweed has a core set of genes with few retained paralogs (expanded families). In contrast, species like Arabidopsis, rice, Brachypodium and Maize have almost 20% of their genes in orthogroups with more than 10 genes (Figure S8). There were 408 and 635 Wolffia and Spirodela specific OGs respectively (as compared to rice, Arabidopsis, Zostera and banana), and 77 OGs exclusive to both (Figure 3A). The OGs specific to Wolffia can be summarized into the significant gene ontology (GO) categories (FDR <0.05) of sphingolipid biosynthesis, photomorphogenesis, wax metabolism and cofactor catabolism (Figure 3B; Table S7). We took a look at the genes that made up these Wolffia specific OGs using our annotation, and we found genes are associated with cell wall architecture (Fasciclin-like arabinogalactan proteins) (Johnson et al. 2011), environment-specific expression orchestration (Nuclear transcription factor Y) (Zhao et al. 2016), flowering time (Casein kinase 1-like HD16) (Hori et al. 2013), and sphingolipids (Huby et al. 2020) (Figure 3D; Table S8).

**Figure 3.**
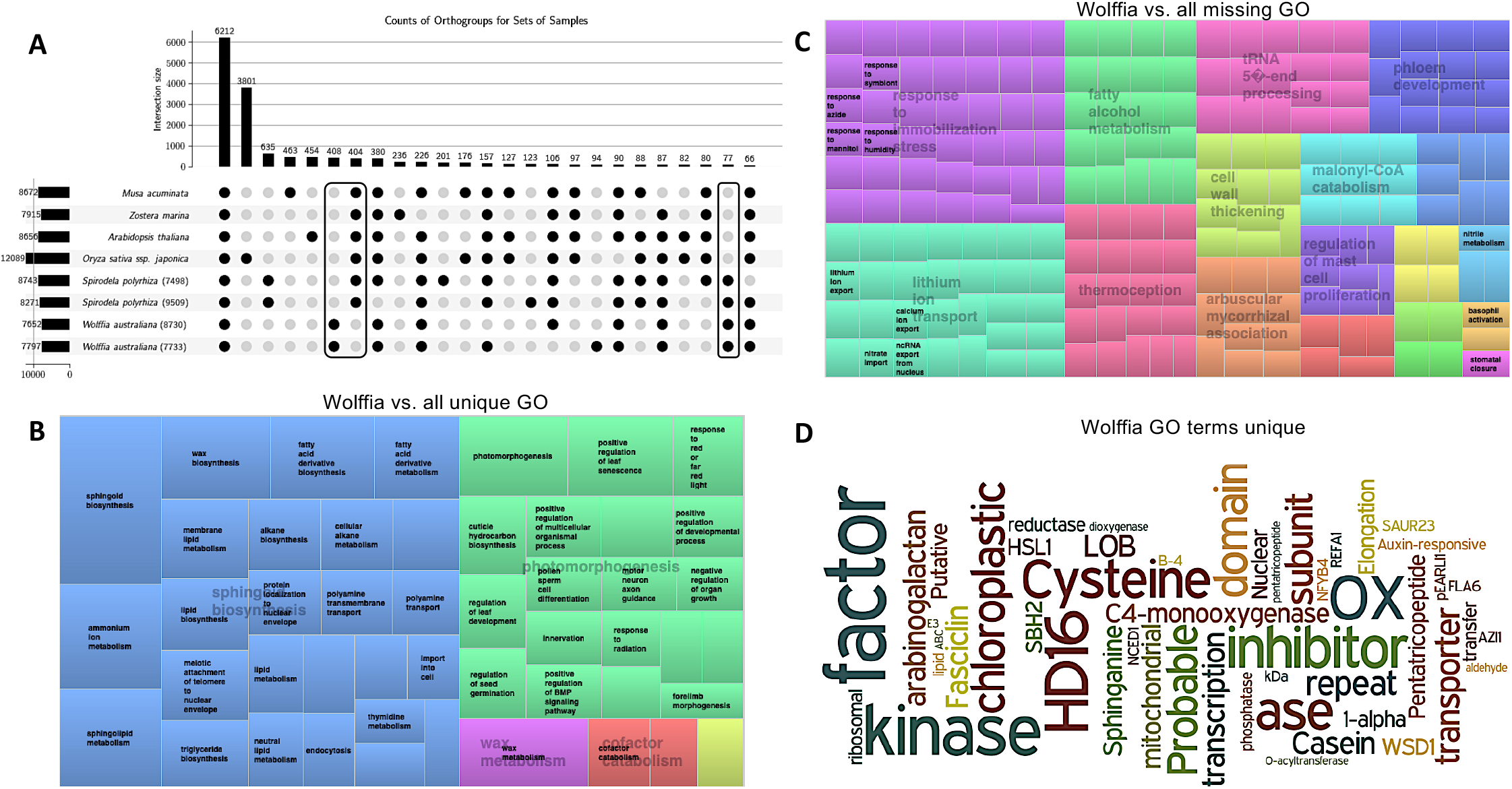
Wolffia genome focused on growth. **A)** An upplot graph showing the orthogroups (OGs) present (black circles) or missing (grey circle) across the four duckweed species (sp9509, sp7498, wa7733 and wa8730), model species (*Arabidopsis thaliana, Oryza sativa ssp. Japonica*), and two non-grass monocots (*Musa acuminate, Zostera marina)*. Boxes indicate Wolffia specific, Wolffia missing and duckweed specific genes. **B)** The OGs that are unique (completely missing in the other species) to both Wolffia genomes were used to identify significantly overrepresented gene ontology (GO) terms (FDR<0.05), summarized using REVIGO, and biological processes were plotted in treemap view. **C)** The OGs that are completely missing in both Wolffia genomes but found in the other species were used to identify significantly overrepresented gene ontology (GO) terms (FDR<0.05), summarized using REVIGO, and biological processes were plotted in treemap view. **D)** The OGs that were unique to both Wolffia genomes were used to identify all genes in those OGs and the gene annotation was used to generate a word cloud to show Wollfia specific terms. Common terms were removed to increase clarity on informational terms.

In contrast, the significant OGs (FDR >0.05) missing in Wolffia included response to immobilization stress, lithium ion transport, fatty alcohol metabolism, thermal perception, cell wall thickening, and phloem development (Figure 3C; Table S9). Looking at specific genes with known function suggests there is a loss of OGs involved in meristem development (*FAF*, *BON1*, *SCL3/11*, *POLAR*, *PSD*, *SZ1*), chromatin (*MED4*, *POLD3*), and light signalling (*BBX12*, *DOT3/NPH3*, *GBF4*) (Table S9). Specifically, *dot3* mutants in Arabidopsis have defects in shoot and primary root growth and produce an aberrant parallel venation pattern in juvenile leaves (Petricka et al. 2008), whereas *nph3* mutants were first identified from genetic screens for Arabidopsis mutants impaired in hypocotyl and root phototropism (Liscum and Briggs 1995). In contrast to Arabidopsis, Spirodela has only one *DOT3/NPH3*, which has also apparently been lost via insertion of LTRs in the Wolffia genomes (Figure 2D). Another very interesting OG that is completely missing in Wolffia but has a large family in Arabidopsis (11), rice (9) and most other land plants is the *CASPARIAN STRIP MEMBRANE DOMAIN PROTEIN* (*CASPs*) Family (Roppolo et al. 2014).

Some of the most significant missing GO terms were in the terpene biosynthesis pathway, while the most significant Wolffia-specific GOs were genes predicted to be in the sphingolipid related pathways. Since both terpenes and sphingolipids may play a predominant role in plant defense (Singh and Sharma 2015; Huby et al. 2020), it is possible that Wolffia has traded the terpene pathway for sphingolipids or that the aquatic environment favors the latter. Related to genes involved in defense, one of the most conserved gene families in plants is the NLR (Nucleotide-binding Leucine-rich Repeat) superfamily encoding hundreds of disease resistance (R) genes that provide innate immunity to pathogen associated molecular patterns or effector triggered resistance responses that are associated with systemic activation of broad spectrum immunity (Jones and Dangl 2006). Members of this family of proteins are known to be the most rapidly evolving genes in plants and they are likely under strong selection pressure (van Wersch and Li 2019). In the sp9509 genome, we have previously annotated only 58 NLR genes in striking contrast to the 178 NLRs that are known in Arabidopsis and 387 that are in Brachypodium (Michael et al. 2017). Remarkably, in our draft genomes for wa7733 and wa8730, we found only a single canonical NLR gene, which also have conserved homologs in the *S. polyrhiza* genomes but is very divergent from any other species. Additionally, two NLR-like genes that contained incomplete NB-ARC domains were identified in the *W. australiana* genome assemblies and these genes are highly conserved across duckweeds, Arabidopsis and rice. The dramatic attrition of this highly conserved gene family in Wolffia is strong evidence that it has further economized its strategy for growth by minimizing this particular type of conserved pathogen defense system. The case of the Wolffia genome demonstrates that the NLR gene family could be largely dispensable for plant survival. It would be intriguing to unravel the mechanism(s) that Wolffia has evolved to cope with the microbial invaders in its nutrient rich aquatic habitat independent of a significant repertoire of NLR immune receptors.

Another surprise was the loss of genes associated with light signaling and the circadian clock in Wolffia. We reasoned that since Wolffia has apparently optimized for fast growth through rapid multiplication that it may have expanded light signaling and circadian gene families in order to better fine-tune its physiological response to the environment. However, we observed the opposite with Wolffia, which only had one third of these genes as compared to that in other land plants (30 vs. ~90), and half to that of Spirodela and Zostera (Table 3). Of the conserved single copy BUSCO genes Wolffia is missing *WITH NO LYSINE (K) KINASE 1* (*WNK1*) (Murakami-Kojima et al. 2002), *TANDEM ZINC KNUCKLE* (TZP) (Loudet et al. 2008), *FAR1-RELATED FACTOR1* (*FRF1*) (Ma and Li 2018) and several of the light harvesting complex proteins. In addition, Wolffia is missing the core clock component *TIMING OF CAB EXPRESSION 1* (*TOC1/PRR1*), which is also missing in Spirodela and Zostera. TOC1/PRR1 specifically binds the promoter of *CELL DIVISION CONTROL 6* (*CDC6*) to regulate the time of DNA replication licensing and growth in Arabidopsis (Fung-Uceda et al. 2018). Many of these genes play multiple roles across signaling, development and stress-response pathways, suggesting Wolffia may have lost these genes for streamlined growth by minimizing cross-talk and checkpoints for feedback control.

**Table 3.**
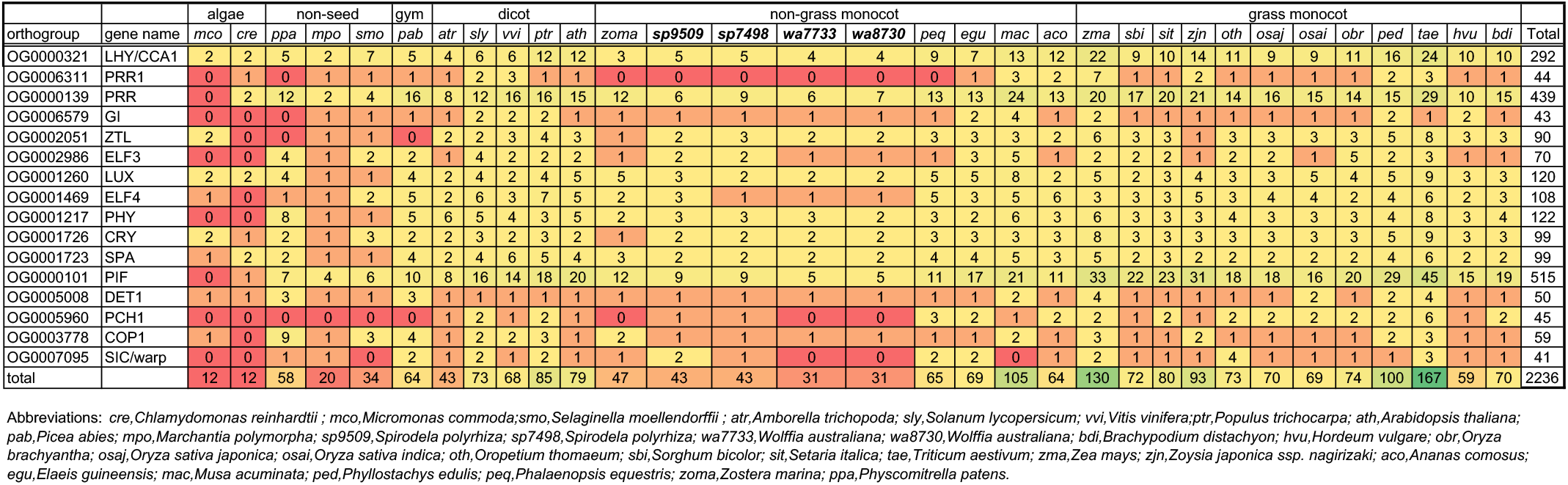
Circadian, light and growth related gene counts across species.

### Time-of-day (TOD) networks

Duckweeds and specifically Wolffia are the fastest growing plants on Earth (Ziegler et al. 2015; Sree et al. 2015b); *W. australiana* doubles in just over a day (Table 1). Since growth is controlled at some level by the circadian clock and coordination of TOD expression networks, we set out to produce a temporally-resolved transcriptome dataset for one of the two Wolffia accessions. Wa8730 was grown under standard photocycles and constant temperature, which is 12 hours (hrs) of light (L) at 20°C, and 12 hrs of dark (D) at 20°C (referred to as LDHH, light/dark/hot/hot) for several weeks. Fronds were then sampled every 4 hrs over 2 days for a total of 13 time points (Figure S9). Total RNA was sequenced and gene expression was estimated based on the predicted gene models. We found that 83% (11,870) of the predicted genes were expressed significantly under the tested conditions (Figure S10; Table S10). We estimated the number of genes showing cycling behavior (R>0.8) and found that 13% (1,638) of the expressed genes displayed a TOD expression pattern (Figure S10; Table S11). This is a smaller percent of cycling genes compared to other plants tested under LDHH, such as Arabidopsis (45%), rice (41%), poplar (30%), douglas-fir (29%) and Brachypodium (27%) (Michael et al. 2008b; Filichkin et al. 2011; MacKinnon et al. 2019; Cronn et al. 2017). Wa8730 genes displayed peak expression, or phase, every hour over the day, similar to LDHH TOD time courses in other plants with more genes peaking at morning or evening specific phases (Figure 4; Figure S11).

**Figure 4.**
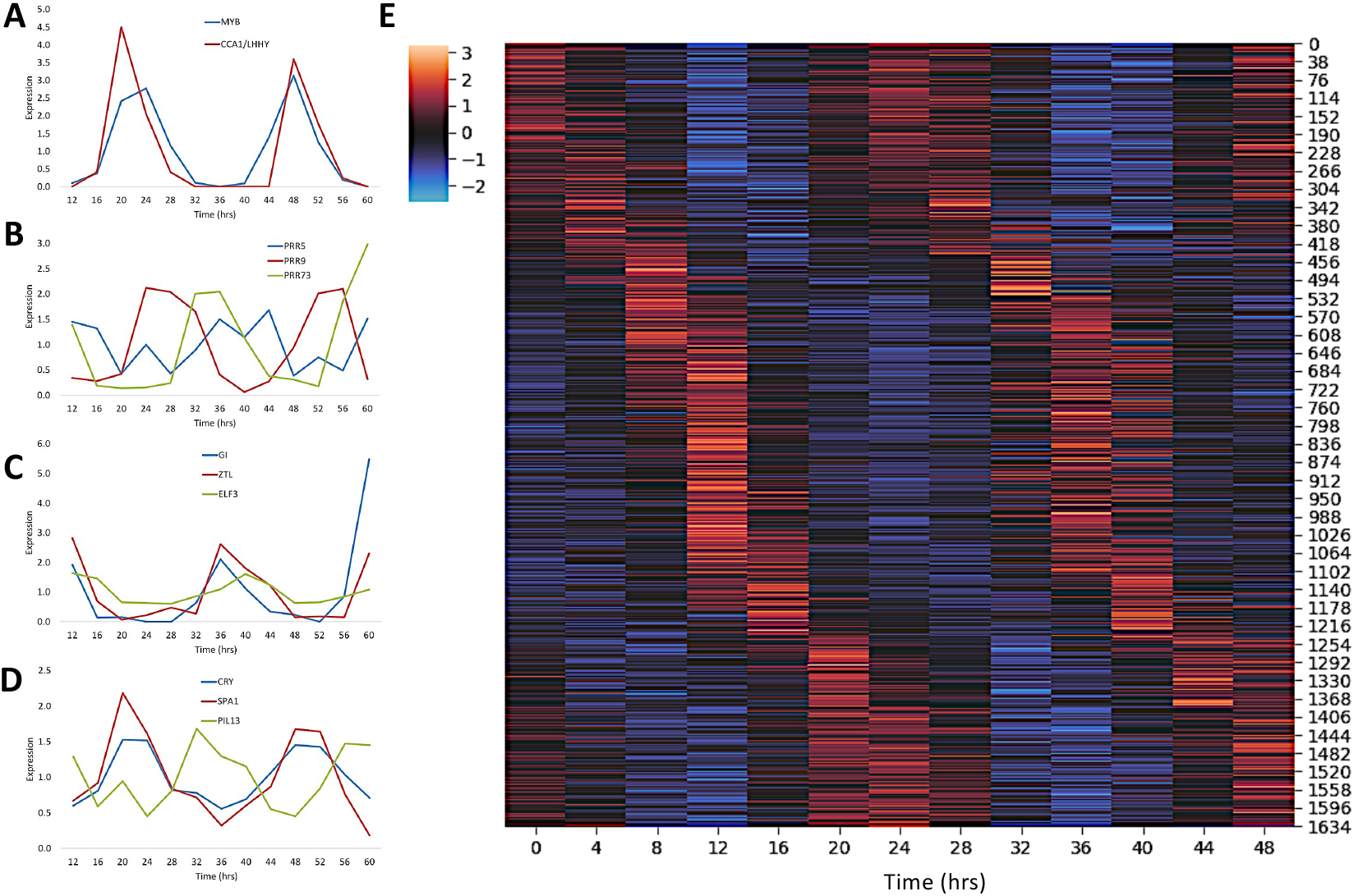
Core circadian and light genes cycle in Wolffia with global expression over the day. **A-D)** Circadian clock and light signaling TOD gene expression in wa8730 is similar to that found in other plants. **A)** MYB (blue) and CCA1/LHY (red) cycle with dawn specific expression. PRR5 (blue), PRR9 (red), and PRR73 (green) show waves of expression similar to other species. **C)** GI (blue), ZTL (red) and ELF3 (green) cycle with evening specific expression. **D)** Light signaling genes CRY (blue), SPA1 (red) and PIL3 (green) cycle over the day. **E)** Heatmap of the cycling genes with red indicating high expression and blue indicating low expression. Genes are on the y-axis and time is on the x-axis.

The phase of expression of core clock genes is conserved across species (Filichkin et al. 2011). Therefore, we looked at the expression of the core genes to both validate the time-course and establish if the expression is also conserved in Wolffia. At the core of the clock is a family of single MYB domain transcription factors LHY and *CIRCADIAN CLOCK ASSOCIATED 1* (*CCA1*), along with a related family called *REVEILLE* (*RVE*) (McClung 2019). While Arabidopsis has 12 single-MYB proteins similar to CCA1/LHY/RVE, Wolffia has 4 and only 2 cycle with a dawn phase; one is the CCA1/LHY ortholog and the other is related to RVE7 (Figure 4A). The other half of the core circadian clock negative-feedback loop is the evening expressed TOC1, which is missing in Wolffia, Spirodela and Zostera. TOC1 is part of a five member gene family of PRRs, which display “waves” of expression across the day (PRR9, dawn; PRR7, midday; PRR5, dusk; PRR3, evening; TOC1/PRR1, evening) (Michael and McClung 2003). Wolffia only had three PRRs, WaPRR9, WaPRR7/3 and WaPRR5, which are phased to dawn, dusk and evening respectively (Figure 4B). Other core circadian genes such as *GIGANTEA* (GI), *EARLY FLOWERING 3* (*ELF3*) and *FLAVIN-BINDING, KELCH REPEAT, F-BOX* (*FKF1*) have evening expression as in other plants (Figure 4C), as well as circadian-regulated light signaling genes (Figure 4D). Despite having a reduced set of circadian and light signaling genes, TOD expression is conserved across the core circadian clock genes.

As another check of the time course, we looked to see if TOD cis-elements that we have found to be conserved in other plant species are also found in Wolffia (Michael et al. 2008b). We searched promoters (500 bp upstream) of genes predicted to be expressed at the same time of day, and found the same cis-elements that we have found across all other plants tested to date (Michael et al. 2008b; Zdepski et al. 2008; Filichkin et al. 2011). For instance, the Evening Element (EE: AAATATCT), which was identified in early microarray experiments and promoter bashing (Harmer et al. 2000; Michael and McClung 2002), is highly overrepresented in the promoters of evening expressed genes (Figure 5A,D), while the Gbox (CACGTG) and its derivatives are highly overrepresented at dawn (Figure 5C), and the Protein Box (PBX: GTGGGCCCC) is overrepresented late in the night (Figure 5E, Table S12,S13). However, the Telobox (TBX: AAACCCT), which is usually overrepresented in genes expressed around midnight, was not significant in Wolffia (Figure 5B). The lack of the TBX could explain at some level the decreased number of cycling genes or it could mean that Wolffia genes with the TBX do not cycle like they do in Arabidopsis, rice and poplar (Filichkin et al. 2011).

**Figure 5.**
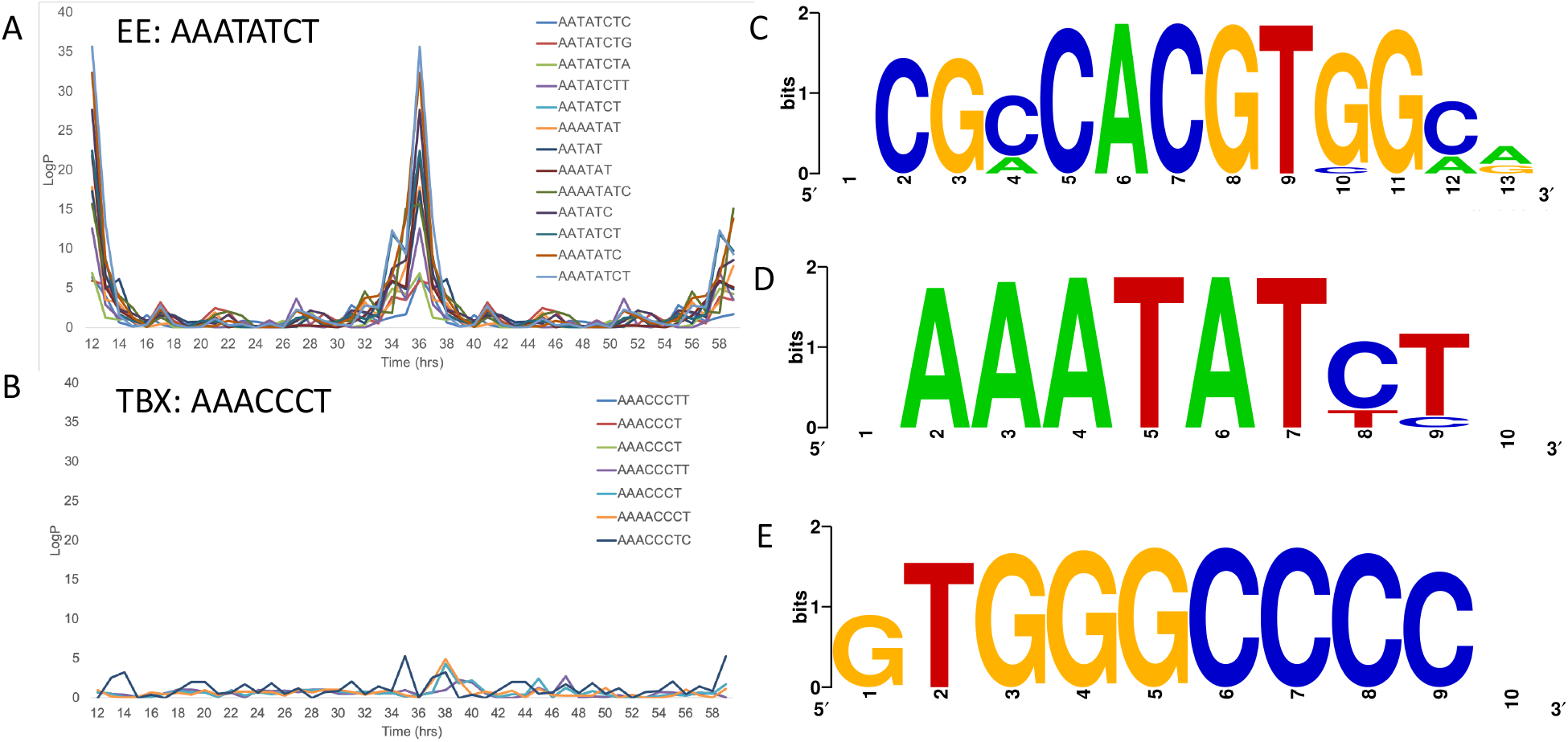
Wolffia has conserved TOD cis-elements but lacks others. **A)** The Evening Element (EE:AAATATCT) is overrepresented in genes with evening-specific expression. **B)** The Telobox (TBX:AAACCCT) is not significantly overrepresented in Wolffia promoters of cycling genes. **C)** Sequence logo of the significantly overrepresented Gbox (CACGTG) in Wolffia. **D)** Sequence logo of the EE overrepresented cis-elements. **E)** Sequence logo of the Protein box (PBX: TGGGCCC) overrepresented cis-elements.

In past studies we hypothesized that if these TOD expression networks are conserved then we should be able to transfer the phase information from one organism to another leveraging its one-to-one (1-2-1) ortholog. We have shown that this works across distantly related plants like Arabidopsis, rice, papaya and poplar both informatically and empirically (Filichkin et al. 2011; Michael et al. 2008b; Zdepski et al. 2008). Therefore, to test whether the TBX was missing in Wolffia or that those genes in this species don’t cycle, we assigned the phase of the Arabidopsis 1-2-1 best ortholog to Wolffia. As an additional control, we also did the same for the picoeukaryote algae *Micromonas commoda* (Mco). We found that most of the known TOD cis-elements were overrepresented in Wolffia and Mco with the EE, TBX and GATA elements highly significant (P<0.05) (Figure 6). Specifically, the TBX was overrepresented in the promoters of Wolffia orthologs assigned the phase from Arabidopsis consistent with Wolffia TBX-containing genes not cycling under the conditions tested or not being rhythmic at all. In Arabidopsis, genes associated with protein synthesis and other activities that occur in the middle of the night contain the TBX cis-element, suggesting Wolffia may have lost the TOD coordination for these pathways.

**Figure 6.**
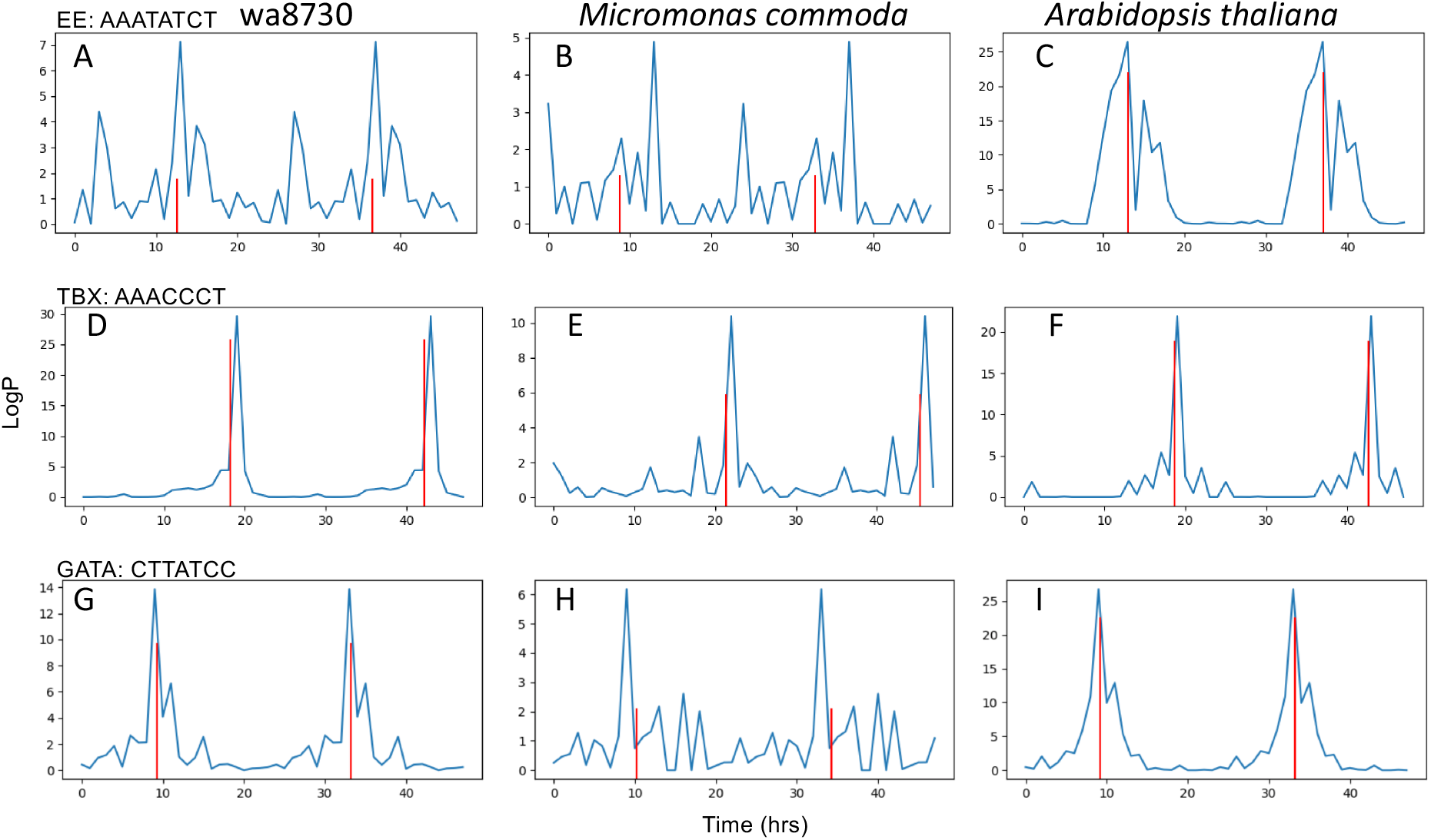
Conserved TOD cis-elements from Algae to Arabidopsis. **A-C)** The EE is overrepresented across all three species. **D-F)** The TBX is overrepresented across all three species. **G-I)** The GATA cis-element is overrepresented across all three species. Arabidopsis cycling genes under LDHH were used to find Wolffia and *Micromonas commode* (Mco) orthologs, and the phase was assigned to those genes. Promoters were then searched for overrepresented cis-elements.

### Wolffia has a focused set of cycling processes

Biological processes are phased to specific times over the day and it is this phasing that controls processes such as growth (Michael et al. 2008a). Since Wolffia has fewer TOD controlled genes and the TBX cis-element was not overrepresented in the promoters of cycling genes like in other plants, we wanted to ask what pathways were no longer under TOD control in Wolffia. Therefore, we compared the Wolffia cycling genes against two high quality datasets generated from Arabidopsis (Ath) and rice (Osa) under the same conditions of light-dark cycles and constant temperature (LDHH) used here (Michael et al. 2008b; Filichkin et al. 2011). In general, cycling genes had similar phases of expression between species as seen in other across-species comparisons (Filichkin et al. 2011; Reyes et al. 2017), although there were some differences in the phase of Wolffia cycling genes (Figure S12). As expected, certain GO terms are overrepresented at specific TOD patterns consistent with Wolffia partitioning its biology over the day much like Ath and Osa (Figure 7A; Figure S13). However, Wolffia only had 92 GO terms significantly overrepresented in a TOD fashion, while Ath and Osa had over double the number of overrepresented GO terms at 238 and 253 respectively (Table S14-16). To look at the reduction in GO terms, we took all cycling genes regardless of TOD expression and found that there were only 7 GO terms shared across all three species, which were all related to photosynthesis (Figure 7B,C; Table S17). This is consistent with the fact that most GO terms overrepresented in Wolffia are related to photosynthesis (Figure 7D). In contrast, Ath and Osa have overrepresented GO terms that span an assortment of biological processes consistent with a global TOD control in these more complex plants (Figure 7E, F) (Michael et al. 2008b; Filichkin et al. 2011). These results could mean that Wolffia has released TOD control over higher order processes, while only coordinating those involved with energy generation. Since the primary role of the circadian clock is to gate specific processes (Michael et al. 2003), and plants can still function without key clock control genes, the reduced TOD control in Wolffia could reflect an unrestricted growth pattern.

**Figure 7.**
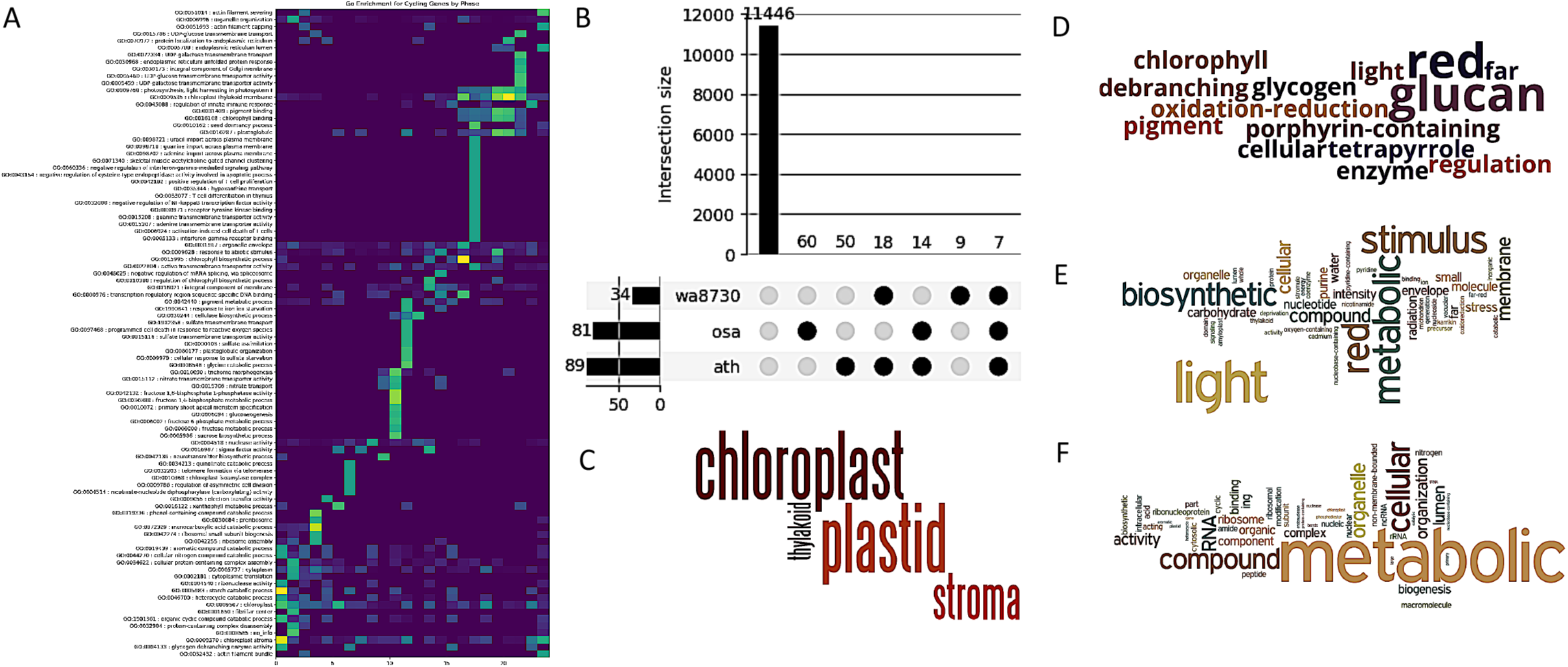
Wolffia TOD control is limited to photosynthesis. **A)** GO term overrepresentation by TOD for Wolffia. **B)** Significantly overrepresented GO term overlaps across Wolffia (wa8730), rice (osa) and Arabidopsis (ath); **C)** Word cloud of the shared significant GO terms across wa8730, osa and ath. **D)** Word cloud of significant GO terms for wa8730. **E)** Word cloud of significant GO terms for ath. **F)** Word cloud of significant GO terms for osa.

## Discussion

Wolffia is the smallest and fastest growing flowering plant on Earth. Here we present two draft genomes for different accessions of *W. australiana*, which has a relatively small genome and contains a minimal set of about 15,000 genes. The loss of genes directly reflects its minimal body plan (i.e. no roots and vasculature) and continuous growth (i.e. fewer clock and development genes), and which were apparently lost through LTR disruption by comparison with Spirodela. A key finding of our work is that Wolffia has a reduced number of TOD regulated genes (13%) compared to other plants, and the genes that remain regulated are specific to photosynthesis and carbon metabolism. Since Wolffia is small (around 1 mm in size), fast growing (DT ~1d), has a minimal set of core plant genes, and grows in direct contact with the media, it offers advantages analogous to the yeast system, which opens up new research opportunities in plants.

*W. australiana* has the smallest genome across the 11 species in the genus tested, which have an average size of 1,136 Mb, and is half the size of the next smallest (*W. brasiliensis* at 776 Mb) (Wang et al. 2011). Despite the range in genome sizes, all Wolffia species have a reduced body plan of a frond with just several thousand cells and no roots (Figure 1; Borisjuk, L., unpublished). They also have a similar fast growth rate of around a day (Table 1) with no obvious relationship to genome size (Sree et al. 2015b). Wolffia is the most derived of all of the duckweed species (Figure 1C), with dramatic body plan changes compared to the most basal Spirodela. The results that several key genes associated with root and light-specific development were disrupted by LTRs in *W. australiana* vs. Spirodela suggests that at some level the changes in morphology are the result of TE activity. Like Spirodela, which also has a small genome, the LTR solo::intact ratio is high in *W. australiana* consistent with LTRs being purged through illegitimate recombination (Devos et al. 2002). Therefore, it is probable that *W. australiana* has a small genome due to the purging of LTRs that in the past have helped to shape its unique gene repertoire and body plan.

Wolffia, and the Lemnaceae family in general, represent extreme examples of plant morphology minimization, and these adaptations are reflected in their reduced yet representative gene sets. Multiple independently assembled genomes of both Spirodela and Wolffia share a common set of missing BUSCO genes (Figure S6), and many of these genes in Wolffia represent genes associated with its morphological innovations (Figure 2; Table S5). Low BUSCO scores have also been noted in other plants with morphological innovations like the parasitic dodder plant (Cuscuta) and carnivorous bladderwort (Utricularia) that both also lack roots and leaf structures like Wolffia (Ibarra-Laclette et al. 2013; Sun et al. 2018; Vogel et al. 2018). *Cuscuta australis* shares many of the same gene losses as Wolffia such as WOX5 (root apical stem cell maintenance), LOP1 (leaf patterning and root development) and the entire family of casparian strip genes (CASP) (Sun et al. 2018). Wolffia is also missing several families of the small signaling peptide CLAVATA3/ESR-RELATED (CLE) that regulate various aspects of cell fate and meristem size (Jun et al. 2010). Despite the extensive gene loss, Wolffia still maintains a core set of gene families (orthogroups, OGs) common to flowering plants (Table S6; Figure S8).

Wolffia has lost a host of genes associated with the intersection between light signaling, phytohormones, circadian clock, growth, immunity and development that may provide insight into its floating ball-shaped morphology. It is critical for a multi-organ plant to position itself relative to the Earth (gravitropism) and the sun (phototropism) for proper development, as well as the means to communicate between the different parts of the organism through systemic signaling (Vandenbrink et al. 2014). Wolffia has lost many genes of the NPH3/RPT2-Like family (NRL),which are required for several auxin-mediated growth processes, including phototropism (root and shoot), petiole positioning, leaf expansion, chloroplast accumulation, stomatal opening and circadian control of PSII photosynthetic efficiency (Christie et al. 2018). NPH3 is the founding member of the NRL family with a close paralog DOT3 that fail to exhibit phototropism and have aberrant venation patterning in Arabidopsis mutants respectively (Petricka et al. 2008; Motchoulski and Liscum 1999). Compared to Arabidopsis, Spirodela only has one NPH3/DOT3, which is lost in Wolffia through a LTR disruption (Figure 2D). In addition, Wolffia is missing the family of LAZY proteins that act as central integrators of gravity sensing with the formation of auxin gradients to control plant architecture (Yoshihara and Spalding 2017). The loss of these two key phototropic and gravitropic pathways provide clues as to how Wolffia has streamlined its body plan.

Loss of the CLE family members, including the CLE3 peptide known to play key roles in orchestrating meristem size in plants is correlated with the unconventional mode of organogenesis in duckweed in which the stipe tissue functions as the site where new meristems are continuously initiated and develop sequentially (Figure 1C) (Lemon and Posluszny 2000). Lastly, the dramatic attrition of the highly conserved NLR family that are known to contain many R genes required to mediate defense signaling and systemic resistance indicates that these genes are largely dispensable for a fast-growing and structurally simple plant. Elucidating how the stipe tissue functions to generate new meristematic centers that are orchestrated to produce new daughters and the mechanism that provides Wolffa with robust defense to pathogens in an R-gene independent manner will reveal much new plant biology. For the latter, the emphasis of the sphingolipid-related pathways observed in our work here may suggest their importance for defense signaling in Wolffia while there are also some evidence for an amplification of the antimicrobial peptide pathways in *S. polyrhiza* (An et al. 2019). These suggestions remain to be tested.

One of the best described growth processes at the molecular level is that of hypocotyl elongation in Arabidopsis (Creux and Harmer 2019). The circadian clock restricts or “gates” the growth during the night hours (dark) through the core clock gene TOC1 binding to the PHYTOCHROME INTERACTING FACTORS (PIF) transcription factors and interaction with one of the core feedback loops mediated by the Evening Complex (EC: ELF3/ELF4/LUX) (Seluzicki et al. 2017). Wolffia, Spirodela and the monocot seagrass *Zostera marina* are all missing TOC1 while having several other PRRs, suggesting that they may replace TOC1 function in aquatic non-grass monocots. In addition, Wolffia is missing the orthologs for PIF3/4 while having the other factors of the EC. Also light regulated growth is controlled at some level by two distinct types of nuclear photobodies, one of which is defined by TANDEM ZINC KNUCKLE/PLUS3 (TZP) (Huang et al. 2016). TZP mutants in Arabidopsis are smaller while overexpression results in large plants that don’t stop growing, consistent with it being a central integrator of light (PHYA) signaling into plant growth (Loudet et al. 2008; Huang et al. 2016; Zhang et al. 2018). The loss of TZP (a BUSCO gene) in Wolffia, and its retention in Spirodela, suggests that a different growth regulation pathway exists or that growth is uncoupled from light in Wolffia. Overall, the loss of growth gating pathways in Wolffia is consistent with the significant decrease in circadian, light and flowering time genes (Table 3), which is in contrast to some CAM plants and crop plants that display expansion of circadian genes (Wai et al. 2019; Lou et al. 2012). These results suggest that the genome innovations responsible for the change in body plan in Wolffia may be closely linked to the loss of specific light-gated growth.

The finding that Wolffia has fewer cycling genes is a bit of a paradox. Ostreococcus is a single-cell alga that is the smallest eukaryote (picoeukaryote) with a 13 Mb genome but a functional circadian clock made up of the core negative feedback loop of CCA1-PPR1 and almost all of the genes cycling in a TOD fashion (Monnier et al. 2010). Similarly, the model microalga Chlamydomonas has a core circadian clock and 80% of its genes cycle in a TOD fashion (Zones et al. 2015). In contrast, multi-cellular, multi-organ plants have around 40% of their transcriptome TOD controlled (Michael et al. 2008b; Filichkin et al. 2011; MacKinnon et al. 2019; Cronn et al. 2017). Therefore, it would seem that a simple (i.e. fewer celled) organism like Wolffia would have almost all of its transcriptome TOD regulated while multi-organ plants would have some processes that would not require TOD expression. The fact that only the core photosynthetic and carbon pathways remain under TOD control suggests that most processes in Wolffia are uncoupled from the environment. A similar situation has been observed in gymnosperms where TOD expression is lost for all but genes associated with photosynthesis in overwintering needles (Cronn et al. 2017). In gymnosperms the hypothesis is that in overwintering needles that are not growing, only core photosynthesis is required. Similarly, a Wolffia frond does not grow very much in terms of cell expansion and most of the growth is due to the rapid cell division associated with the daughter frond in the very limited pocket area of the frond (Figure 1C).

Since Wolffia is in direct contact with the environment (water) where temperature and nutrients are most likely in a more or less constant state over the day, it is possible that Wolffia has uncoupled these processes from TOD expression required in an environment on land. While most conserved TOD cis-elements are found (Figure 5) (Michael et al. 2008b), the loss of TOD overrepresentation of the TBX in Wolffia, but yet the identification of it through cycling-orthologs with Arabidopsis (Figure 6), suggest that the highly conserved TBX-controlled pathway are not TOD regulated in Wolffia (Filichkin et al. 2011). The loss of TBX regulation could reflect a general loss of key regulatory switches (TFs and others) associated with the circadian clock and light signaling (Table 3). Since many of the gene losses in Wolffia link development with light signaling, it is possible that the evolutionary path to a highly reduced plant with simple architecture and continuous growth also resulted in the loss of light-specific gated growth.

Wolffia is like the yeast of flowering plants with a core set of angiosperm genes, small size, rapid unrestricted growth, and growing in direct contact with its environment. Before Arabidopsis, duckweed was widely used as a model plant in Plant Biology (Lam et al. 2014). In fact, photoperiodic flowering was first worked out in duckweed (Hillman 1976) while the auxin biosynthetic pathways were characterized by radioisotope labeling in *Lemna gibba* (Rapparini et al. 1999). Wolffia still has the genes for flowering and could be developed as a genetic system that is distinct from that of Arabidopsis, where crosses may be made by mixing flowering strains and collecting the seeds that sink to the bottom. The limited number of cells and cell types in Wolffia provide a simplified model to dissect cell specific regulation and how plant cells directly respond to specific chemicals at the organismal level.

## Material and Methods

### Wolffia growth rate experiments

The four *W. australiana* (Benth.) Hartog & Plas accessions 7211 (Australia, victoria), 7540 (New Zealand), 7733 (Australia, South Australia), and 8730 (Australia, New South Wales) were maintained at the Rutgers Duckweed Cooperative (http://www.ruduckweed.org/) or at the stock collection at the University of Jena, Germany. All four *W. australiana* accessions were pre-cultivated for four weeks with a subculture interval of 1 week at 25°C and continuous white light of an intensity of 100 μmol m^−2^s^−1^. For growth rate measurement, 10 ± 1 fronds from the pre-cultures were inoculated into 100 ml nutrient medium N contained in 150 ml beakers (Appenroth et al. 1996). The plants were grown for 1 week at 25°C and continuous white light of an intensity of 100 μmol m^−2^s^−1^. The final frond count was taken on day 7. The experiment was carried out with 6 replicates and was repeated at an interval of 1 month. All values are average ±SE. Explanation of Relative Growth Rate (RGR), Doubling Time (DT) and Relative (weekly) Yield (RY) can be found in Ziegler et al., (Ziegler et al. 2015).

### Ultrastructure of *W. australiana* and Imaging

*W. australiana* was grown on 2% agar (Agar‐Agar, danish, Carl Roth GmbH, Karlsruhe, Germany) with half-strength MS salts (Murashige and Skoog 1962) under sterile conditions at 25 ± 1°C (white light of 85 μmol m^−2^s^−1^ 16 h day and 8 h night photoperiod). Frond proliferation was monitored using a digital microscope Keyence VHX-5000 (Keyence Deutschland GmbH, Neu-Isenburg) by capturing images at 30 min intervals during 45 hours. In order to demonstrate growth dynamic the video file was converted to time‐lapse movie MP4 format.

For histological examinations, intact plants were fixed in FAA (Weigel and Glazebrook 2008), infiltrated with Spurr Resin (Plano, Wetzlar, Germany) and polymerized at 70°C for 24 h. The serial block sectioning was performed using Semimikrotom Leica Ultracut UTC (Leica Mikrosysteme GmbH, Vienna, Austria); the semi-thin 1 μm tissue sections were stained with Methylenblau-Borax (Waldeck GmbH & Co. KG, Muenster) and the stack of 311 micro images was obtained using digital microscope Keyence VHX-5000 with 600× lens magnification (Keyence Deutschland GmbH, Neu-Isenburg) and Image J software (Schindelin et al. 2012). The creation of three-dimensional models from a set of images (3D-reconstruction) was performed using Amira-Avizo software (ThermoFisherScientific).

### TOD time course growth, RNA extraction and sequencing

For the 2-day time course experiment, roughly 600 mg of tissue was sampled every 4 hr over 2 days for a total of 13 time points for wa8730 and at 0 hr and 12 hr for a total of 2 time points for wa7733 (Figure S9). Samples were collected in the dark under a dim green light, flash frozen in liquid nitrogen and stored at −80 °C. RNA was extracted using the total RNA isolation protocol from mirVana miRNA Isolation Kit (Thermo Scientific, Cat. No. AM1560). The RNA quality was validated using gel electrophoresis and the Qubit RNA IQ Assay (ThermoFisher). Stranded RNAseq libraries were constructed using 2ug of total RNA quantified using the Qubit RNA HS assay kit (Invitrogen, USA) with the Illumina TruSeq Stranded Total RNA with Ribo-Zero Plant (Illumina, USA). Multiplexed libraries were pooled and sequenced on an Illumina HiSeq2500 under paired-end 150 bp mode.

### DNA isolation, library construction, and sequencing

HMW genomic DNA was isolated from young teff leaf tissue for both PacBio and Illumina sequencing. A modified nuclei preparation was used to extract HMW gDNA and residual contaminants were removed using phenol chloroform purification (Lutz et al. 2011). The DNA sample was run overnight on a low concentration agarose gel (<0.5% W/V) to ensure the DNA has high molecular weight (>40 kb) with Lambda DNA used as a control. PacBio libraries were constructed using the manufacturer’s protocol and were size selected for 30 kb fragments on the BluePippin system (Sage Science) followed by subsequent purification using AMPure XP beads (Beckman Coulter). The PacBio libraries were sequenced on a PacBio RSII system with P6C4 chemistry. In total, 16 and 36 Gb of filtered PacBio reads were generated, which represents 41× and 91× coverage for wa8730 and wa7733 respectively. The same batch of HMW genomic DNA was used to construct Illumina DNAseq libraries for correcting residual errors in the PacBio assembly. Libraries were constructed using the KAPA HyperPrep Kit (Kapa Biosystems) followed by sequencing on an Illumina HiSeq2500 with 2×150 bp paired-ends (wa7733 and wa7540). Wa8730 was sequenced on an Illumina MiSeq with 2×311 bp paired-ends.

### Genome assembly

PacBio data was error corrected and assembled using Canu (v1.5) (Koren et al. 2017), which produced accurate and contiguous assembly for homozygous plant genomes. The following parameters were modified: minReadLength=2000, GenomeSize=375Mb, minOverlapLength=1000. Assembly graphs were visualized after each iteration of Canu in Bandage (Wick et al. 2015) to assess complexities related to repetitive elements and homoeologous regions. A consensus was first generated using the PacBio reads and three rounds of racon (v1.3.1) (Vaser et al. 2017). The raw PacBio contigs were polished to remove residual errors with Pilon (v1.22) (Walker et al. 2014) using greater than 30x coverage of Illumina paired-end 150 bp data for wa8730 and wa7733 respectively. Illumina reads were quality-trimmed using Trimmomatic followed by aligning to the assembly with bowtie2 (v2.3.0) (Langmead and Salzberg 2013) under default parameters. Parameters for Pilon were modified as follows: --flank 7, --K 49, and --mindepth 15. Pilon was run recursively three times using the modified corrected assembly after each round. The statistics for the final Canu based PacBio assembly are summarized in Table 2. Completeness of the genome was checked by both looking at the mapping of the illumina reads to the genome assemblies, as well as mapping high quality Sanger sequenced cDNA clones. The average Illumina read mapping to the genome assembly at each pilon step was 95 and 97% for wa8730 and wa7733 respectively, consistent with the assemblies covering almost all of the genome space. In addition, 1,987 high quality Sanger sequenced cDNA clones from the Waksman Student Scholars Program (WSSP) (https://www.ncbi.nlm.nih.gov/nuccore/?term=Waksman+Student+Scholars+program+wolffia+australiana) were mapped to the genome assemblies with 100 and 99.7% of the cDNAs mapping to the wa7733 and wa8730 genomes, respectively. The WSSP cDNA sequences were mapped to the wa8730 and wa7733 genome assemblies using minimap2 (v2.17-r941) (Li 2018).

### BioNano Genomics maps

Bionano optical maps were prepared as previously described (Kawakatsu et al. 2016) with minor modifications; High-molecular weight (HMW) DNA was extracted from up to 10◻g whole plant tissue. Briefly, extracted HMW DNA was nicked with the enzyme Nt.BspQI (NEB, Beverly, MA, USA), fluorescently labeled, repaired, and stained overnight according to the Bionano Genomics nick-labeling protocol. Nick-labeled DNA was run on a single flow cell on the Irys platform (Bionano Genomics, San Diego, CA, USA). The IrysView software (Bionano Genomics; version 2.5.1) was used to quality filter the raw data (>100 kb length, >2.5 signal/noise ratio) and molecules were assembled into contigs using the “small optArguments” parameters. Resulting Bionano cmaps were compared against the different assemblies using Bionano RefAlign, and collapsed regions or artificial expansions were detected as structural variations using the structomeIndel.py script (https://github.com/RyanONeil/structome).

### BUSCO single-copy ortholog analysis

We benchmarked each duckweed assembly (wa8730, wa7733, sp9509, sp7498) using the BUSCO (v3) liliopsida odb10 database (Simão et al. 2015). This produces a list of BUSCO genes found to be complete, missing, duplicated or partially found in each genome and forms the basis of a contingency table represented by an upset plot (Table 2; Table S4; Table S5; Figure S6). A chi-square test found that the missing genes in the duckweeds were not independent. Subsequent post hoc tests showed that genes found to be missing in any duckweed were at increased chance of being found missing in the other duckweeds. Further, genes found to be missing in either *Wolffia* were at increased chance of being found missing in the other. (Figure S6). In order to get more information about the missing busco genes, the representative ancestral proteins were blasted against Arabidopsis. Results were manually filtered in order to find interesting missing genes such as WOX5, TPL2, etc.

### Repeat analysis

Several different types of repeats were identified in the Wolffia genomes including long terminal repeat retrotransposons (LTRs) and ribosomal DNA (rDNA) arrays. LTRharvest (GenomeTools v1.5.10) with the specific settings (-xdrop 37 -motif tgca -motifmis 1 -minlenltr 100 -maxlenltr 3000 -mintsd 2) were used to find full length LTRs (Ellinghaus et al. 2008). The full length LTRs, 2,510 and 1,892 in wa7733 and wa8730 respectively, were used as the library to mask their respective genome assemblies with RepeatMasker (v4.0.7) with default settings (Smit et al. 2015). Wolffia rDNA was identified with the Spirodela 18S, 5.8S, 26S and 5S sequence using BLAST against the genome assemblies (Table S3). The Wolffia consensus rDNA sequence was used to search the genomes to determine how many repeats were in each genome, and consistent with the contiguity statistics for two genomes wa7733 has about double the number of assembled rDNA as compared to wa8730 (Table S3). To estimate the number of rDNA repeats in the two Wolffia accessions we performed a coverage analysis with the Illumina short reads (Hoang et al. 2018). We created a library with one full length rDNA array, one 5S array, and the single copy gene GIGANTEA (GI), and mapped the Illumina reads using minimap2 v2.17-r941) (Li 2018). The number of rDNA repeats was estimated by dividing the coverage across the rDNA arrays by the coverage over GI.

### Genome size estimation

The genome size of *W. australiana* has been reported to be 375 +/− 25 Mb and 357 +/− 8 Mb for accessions wa8730 and wa7733 respectively by flow cytometry (Wang et al. 2011). More recently the genome size of wa7540 was estimated by flow cytometry to be 432 +/− 6 Mb (Hoang et al. 2019). These estimates were carried out with different internal standards, which could reflect the differences in size or they could represent real variation between *W. australiana* accessions. We estimated the genome sizes of all three accessions by kmer frequency using Illumina sequencing reads. Using an estimated 68, 80 and 62 fold Illumina sequencing coverage (375 Mb genome size) for wa8730, wa7733 and wa7540 respectively, the kmer (k=19) count was generated using jellyfish (v2.2.4) using no kmer cut off (no -U), which enables counting of up to 10,000 kmers to capture the high copy number repeats (centromere, rDNA and young TEs). The kmer frequency was estimated by analyzing the Jellyfish histo file with GenomeScope (Vurture et al. 2017) and verified with an in-house pipeline (Figure S1). The kmer profiles are consistent with diploid homozygous plants with one predominant peak with heterozygosity ranging from 0.29-0.60%, 31-33% repeat sequence and very similar genome sizes between 342-345 Mb (Figure S1).

The kmer genome size estimates results are consistent with the 357-375 Mb flow cytometry genome size estimate (Wang et al. 2011), but the consistency (within 3 Mb) in kmer genome size estimates suggested the past flow cytometry estimates may not reflect the true genome size variation between accessions. Therefore, we measured all three accessions using the same method as Hoang et al., (Hoang et al. 2019), which uses radish (*Raphanus sativus*) as the reference, and found that the genome size estimates were also similar to one another but higher at 432, 441 and 432 Mb for wa7733, wa8730 and wa7540. Since the genome assemblies sizes were 395 and 355 Mb for wa7733 and wa8730 respectively, and both Illumina and PacBio reads covered greater than 95% of the genome assemblies, which confirms that 100 Mb is not missing from the assemblies, the flow cytometry results most likely are a high estimate of genome size. However, it does confirm that the genome size variation between the accessions is small (1-3 Mb). The differences in assembled genome size (395 vs. 355 Mb) could just reflect the amount of high copy number repeats (centromere and rDNA) (Figure S2; Table S3) assembled based on their coverage (91x vs. 41x), which would be consistent with the kmer genome size estimate of 342-345 Mb (Figure S1).

### Gene and repeat predictions and orthogroup mappings

Custom repeat libraries were constructed for each species following the MAKER-P basic protocol (Campbell et al. 2014). RepeatModeler 1.0.8 was run against the genome assembly to produce an initial *de novo* library (Smit and Hubley 2008-2015). Sequences with blastx hits (e-value 1 × 10^−10^) to a UniProt database of plant protein coding genes were removed along with 50 bp flanking sequences. The resulting custom library was used with RepeatMasker to produce hard- and soft-masked versions of the genome assemblies.

Protein coding genes were annotated for all four duckweed genomes with the MAKER 3.01.02 pipeline (Holt and Yandell 2011). For both sp9509 and wa8730, Illumina RNA-seq reads were trimmed with skewer 0.2.2 (Jiang et al. 2014), aligned to the genome assembly with HISAT2 2.1.0 (Kim et al. 2015), and assembled with Trinity 2.6.6 (Grabherr et al. 2011; Haas et al. 2013) in genome-guided mode. Trinity was run independently in *de novo* mode, and all transcripts from the Trinity runs were aligned to the genome and reassembled with PASA (Haas et al. 2003) to form a comprehensive transcriptome assembly for each species. EST sequences from sp7498 (Wang et al. 2014) were downloaded from SRA (SRR497624) and assembled with Newbler 3.0 (Margulies et al. 2005). All available assembled duckweed transcripts were passed to MAKER as evidence. Protein homology evidence consisted of all UniProtKB/Swiss-Prot (Schneider et al. 2009; UniProt Consortium 2019) plant proteins and the following proteomes: *Arabidopsis thaliana*, *Elaeis guineensis*, *Musa acuminata*, *Oryza sativa*, *Spirodela polyrhiza*, and *Zostera marina (Goodstein et al. 2012; Singh et al. 2013)*.

Three different approaches were used for *ab initio* gene prediction. RNA-seq alignments along with the soft-masked assembly were passed to BRAKER 2.1.0 to train species-specific parameters for Augustus 3.3.3 and produce the first set of coding gene predictions (Stanke et al. 2008, 2006; Hoff et al. 2016, 2019). A second set of predictions was produced by generating a whole-genome multiple sequence alignment of all duckweed species with Cactus (Paten et al. 2011) and running Augustus in CGP mode (König et al. 2016). The final set of predictions was produced by running Augustus within the MAKER pipeline, using the BRAKER-generated species parameters along with protein and assembled RNA-seq alignments passed as evidence.

Finally, MAKER was run and allowed to select the gene models most concordant with the evidence among these three prediction datasets. MAKER-P (Campbell et al. 2014) standard gene builds were generated by running InterProScan 5.30-69 (Jones et al. 2014) and retaining only those predictions with a Pfam domain or having evidence support (AED score < 1.0).

Orthogroups and orthologues were identified across 29 proteomes from the Plaza 4.0 Monocots database (Van Bel et al. 2018) and the duckweed MAKER-P standard build proteomes with alternate transcripts removed. OrthoFinder 2.2.7 (Emms and Kelly 2019, 2015) was run against all-vs-all proteome alignments computed with DIAMOND blastp (Buchfink et al. 2015) using the standard workflow.

### Gene Content Analysis via Orthogroups (OG) and Gene Ontology (GO)

Orthogroup (OG) counts (described above) were used as the basis for MDS in order to visualize relationships among the 29 Plaza monocot (v4) proteomes and the duckweed proteomes. These same counts were processed using Fisher exact tests to identify OG groups for which duckweeds/Wolffia had disproportionately high or low membership. After an FDR correction, we were able to identify OG that were significantly under/overrepresented in duckweeds/Wolffia. In parallel, the 29 Plaza monocot (v4) proteomes and the four duckweed proteomes were processed by eggNOG-mapper (Huerta-Cepas et al. 2017) to create a consistent set of GO term classifications for every protein in each species in the dataset. Frequencies of GO terms for each species were then processed in a similar way to the OG in order to identify GO terms that were significantly under/overrepresented among the duckweeds/Wolffia. The significant GO terms (FDR < 0.05) were summarized and visualized using REVIGO (Supek et al. 2011).

### Annotation of NLRs (Nucleotide-binding Leucine-rich Repeat proteins)

The NLRs were predicted using NLR-Annotator (Steuernagel et al. 2020) that scans genomic sequences for MEME based sequence motifs (Bailey et al. 2009). In addition the proteomes were queried for the presence of the NB-ARC domain (PF00931) (Sarris et al. 2016). Genomic and proteomic NLR predictions were combined to create a non-redundant list of putative NLRs, each putative NLR with an available gene model was then run through interpro-scan (Jones et al. 2014) for further domain prediction. The set of NLRs from either accession and species were then aligned by MAFFT (Katoh and Standley 2016) to identify orthologs.

### RNAseq processing and TOD expression prediction

TOD time course data was analyzed with similar methods that have been described with some modifications described below (Michael et al. 2008b; Wai et al. 2019; MacKinnon et al. 2019; Filichkin et al. 2011). HISAT2 (Kim et al. 2015) was used to align RNAseq reads to the wa8730 assembly. The resultant alignments were processed by Cuffquant and CuffNorm (Trapnell et al. 2012) to generate normalized expression counts for each gene for each time point. Genes with mean expression across the 13 timepoints below 1 FPKM were filtered prior to being processed by HAYSTACK (Michael et al. 2008b) to identify genes that show cycling expression behavior (Figure S10; Table S10). HAYSTACK operates by correlating the observed expression levels of each gene with a variety of user specified models that represent archetypal cycling behavior. We used a model file containing sinusoid, spiking traces, and various rough linear interpolations of sinusoids with periods ranging from 20 hours to 28 hours in one hour increments and phases ranging from 0-23 hours in one hour increments. Genes that correlated with their best fit model at a threshold of R > 0.8 were classified as cyclers with phase and period defined by the best fit model (Figure 4; Figure S11). This threshold for calling cycling genes is based on previous observations (Michael et al. 2008b; Wai et al. 2019; MacKinnon et al. 2019; Filichkin et al. 2011), although in some cases it is too stringent as evidenced by the exclusion of one of the core circadian clock genes LHY that clearly cycles (Figure 4A) but has an R (0.78) just below the threshold. As a check cycling genes were also predicted using JTKcycle (Hughes et al. 2010), and the results were consistent with HAYSTACK and the 0.8 threshold (data not shown). The HAYSTACK cycling classifications formed the basis of our promoter analysis of wa8730 and our GO term analysis of wa8730.

We leveraged two high quality TOD datasets in two well studied model plants *Arabidopsis thaliana* (Michael et al. 2008b) and *Oryza sativa* (Filichkin et al. 2011) to determine how the Wolffia TOD expression compares to other plants. For each orthogroup (OG), for each species, we computed a circular mean of the phases of the cycling genes belonging to that OG in that species. This allows us to use the OGs to map from some cycling gene in wa8730 to a ‘cycling behavior’ in orthologous genes in osa and ath. We used this mapping to create a KDE histogram plot representing the density of these pairs: phase of cycling gene, orthologous ‘cycling behavior.’ This plot represents how cycling behavior in wa8730 genes related to cycling behavior in osa and ath orthologues. For example, genes that cycle and peak at dusk (11-14h) in wa8730 are associated more with dusk genes in osa and ath than with morning genes (Figure S12).

Once cycling genes in wa8730 were identified, we were able to find putative cis-acting elements associated with TOD expression. Promoters, defined as 500 bp upstream of genes, were extracted for each gene in wa8730 and processed by ELEMENT (Michael et al. 2008a, 2008b; Mockler et al. 2007) to compute background statistics. Promoters for cycling genes were split according to their associated phase and kmers that were overrepresented in any of these 24 promoter sets were identified by ELEMENT. By splitting up cycling genes according to their associated phase, we gain the power to identify kmers associated with TOD-specific cycling behavior at every hour over the day (Table S12). Our threshold for identifying a kmer as being associated with cycling was an FDR less than 0.05 in at least one of the comparisons. The significant kmers were manually clustered according to sequence similarity identifying the ME, Gbox, and EE motifs (Table S13). Interestingly, the TBX motif was not identified as significant (Figure 5).

Due to the conserved nature of TOD expression networks across plants, the orthologues across species have similar expression patterns and underlying cis-elemnts in their promoters. We have leveraged the conserved nature of TOD expression networks to predict cycling behavior and cis-elements across phylogenetically diverse plants (Michael et al. 2008b; Zdepski et al. 2008), and validated these predictions with empirical TOD expression data (Filichkin et al. 2011). These predictive tools also enable us to evaluate why genes are not cycling or cis-elements are not significant, such as the case in Wolffia. Therefore, we leveraged a high quality *Arabidopsis thaliana* LDHH dataset (Michael et al. 2008b) to assign the phase of expression to reciprocal best blast (RBB) matches of wa8730 and the green alga *Micromonas commoda* (Mco). The promoters of Arabidopsis, and Arabidopsis phase assigned wa8730 and Mco were analyzed using ELEMENT as above. The significant kmers were evaluated for known elements and TBX was significant across all three species as well as other known elements such as the EE and GATA (Figure 6).

We used GO annotations to try to identify what cellular functions were associated with cycling genes. EggNOG-mapper (Huerta-Cepas et al. 2017) was used to provide annotations for each gene establishing a mapping between GO terms and sets of genes associated with those terms. In total, 6,874 genes in wa8730 could be associated with one or more GO terms from a population of 6,945 GO terms. Cycling calls from HAYSTACK were used to separate genes into 25 categories, one for each phase and non-rhythmic. The elim method was used to identify GO terms that were overrepresented among cycling categories. This allows us to associate GO terms with specific cycling behavior (Figure 7; Figure S13; Table S14-17).

## Data Availability

The raw PacBio data, Illumina resequencing, and RNAseq reads were deposited to the National Center for Biotechnology Information (NCBI) Short Read Archive (SRA) under bioproject PRJNA615235 (Table S18). The genome assembly wa8730 and wa7733 are available from CoGe (https://genomevolution.org/) under genome id56605 and id56606 respectively.

## Author contributions

TPM and EL conceived the study, oversaw the experiments and wrote the manuscript; DB and TCM assembled the genome; FJ, JPS and JRE generated Bionano optical maps and scaffolded the genome; KSS and KJA conducted growth experiments; KJA and JF performed genome size estimations with flow cytometry; SO and LB performed sectioning, microscopy and developed images and time-lapse video of Wolffia; PC and SG carried out tissue sampling and RNA isolation for the circadian transcriptome study; EE and RM predicted protein coding genes and protein family (orthogroup) analysis; EB and KK performed NB-ARC gene curation and analysis; TPM and NH conducted orthogroups, gene ontology terms and TOD analyses.

## Acknowledgements

This material is based upon work supported by the U.S. Department of Energy, Office of Science, Office of Biological & Environmental Research program under Award Number DE-SC0018244. Duckweed research at the Lam Lab is also supported in part by a grant from Hatch project (#12116) from the New Jersey Agricultural Experiment Station at Rutgers University. RM and JRE are Investigators of the Howard Hughes Medical Institute.

## Supplemental information

### Supplemental Figures

**Figure S1. *Wolffia australiana* under growth assay**.

**Figure S2. Genome size estimated by Kmer (k=19) frequency**. **A)** wa7733, **B)** wa8740, and **C)** wa7540. Kmers (19 bp) were counted with jellyfish and the resulting “histo” file was plotted using GenomeScope. **D)** Table of GenomeScope results. All three genomes are estimated to be ~340 Mb with 0.3-0.4% heterozygosity and 100 Mb (30%) repeat content.

**Figure S3. Wolffia is colinear with sp9509**. **A)** wa7733 vs. sp9509 and **B)** wa8730 vs. sp9509. **C)** Summary of the gene level syntenic blocks. Genomes were aligned using the last and syntenic regions based on proteins were identified and plotted using SynMap on CoGe (https://genomevolution.org/coge/).

**Figure S4. Wolffia genomes differ mostly by small INDELs**. **w** a8730 and wa7733 were compared using mummer and differences were plotted using assemblytics.

**Figure S5. Wolffia genome is colinear with sp9509**. Green boxes are forward genes and blue boxes are reverse genes. Grey bars are syntenic regions.

**Figure S6. Wolffia and Spirodela genomes are missing a similar set of BUSCO genes**. An upplot graph showing the number of BUSCO genes missing across the four duckweed genome assemblies: sp9509, sp7498, wa8730, and wa7733. There are 2,174 BUSCO genes that are found in all four duckweed genomes. There are 263 missing BUSCO genes shared across the duckweed genomes.

**Figure S7. Multi-dimensional scaling (MDS) based on orthogroups (OGs) reveals relationships between the grass monocots, non-grass monocots and duckweeds**. Circles highlight the grass, non-grass and duckweed groupings. Monocots (blue): non-grass (circle), *Musa acuminate*, *Phalaenopsis equestris*, *Ananas comosus*, *Elaeis guineensis*; grass (diamond), *Phyllostachys edulis*, *Brachypodium distachyon, Hordeum vulgare, Oryza brachyantha, Oryza sativa japonica, Oryza sativa indica, Oropetium thomaeum, Sorghum bicolor, Setaria italica, Triticum aestivum, Zea mays, Zoysia japonica ssp. nagirizakizoma*; Duckweed (square), wa8730, wa7733, sp9509, sp7498; seagrass (x) *Zostera marina*. Dicot (red): weed (plus), *Arabidopsis thaliana*; basal plant (circle) *Amborella trichopoda*; tree (diamond), *Populus trichocarpa*; crop (square) *Solanum lycopersicum, Vitis vinifera*. Algae (green, x), *Micromonas commode*, *Chlamydomonas reinhardtii*. Non-seed plants, Liverwort (purple, plus), *Marchantia polymorpha*; moss (blue, cross), *Physcomitrella patens*; bryophytes (red, plus), *Selaginella moellendorffii*. Gymnosperms (orange, diamond), *Picea abies*.

**Figure S8. Wolffia and Spirodela genes are found in small gene families (orthogroups)**. 33-41% of Wolffia (wa7733 and wa8730) and Spirodela (sp9509 and sp7498) genes are found in orthogroups (OGs) with only one gene, whereas Arabidopsis, rice, brachy (*Brachypodium distachyon*) and maize have between 12-18% and over 20% in OGs with more than 10 genes. In contrast, less than 10% of duckweed genes are found in OGs with more than 10 genes.

**Figure S9. Wolffia time course design**. Two chambers were set to 12 hrs of light and 12 hrs of dark at 20°C, but 12 hrs apart in opposite phases so plants could be collected over a 12 hr period without collecting during “actual night.” Samples were collected every four (4) hours over two days for a total of 13 time points. Zeitgeber time (ZT) is the time relative to the light dark cycle where ZT0 equals lights on.

**Figure S10. Break down of expressed and cycling genes in wa8730**. Total number of genes reflects the predicted protein coding genes in wa8730. Genes with mean expression across the 13 timepoints below 1 FPKM were considered “not expressed” (2,343, orange). Genes that correlated with their best fit model at a threshold of R > 0.8 were classified as “cycling” with phase and period defined by the best fit model (1,638; 13%). The rest were classified as “expressed not cycling” (10,232).

**Figure S11. Distribution of cycling genes over the day**. **A)** wa8730, **B)** Rice (*Oryza sativa*), **C)** Arabidopsis (*Arabidopsis thaliana*). Genes that correlated with their best fit model at a threshold of R > 0.8 were classified as “cycling” with phase and period defined by the best fit model.

**Figure S12. Comparison of cycling genes between Wolffia (wa8730), Arabidopsis (ath) and rice (osa).** Predicted phases were compared between wa8730 and osa **(A)**, as well as wa8730 and ath **(B)**. Density (darker blue) represents the number of genes with an associated phase. Using the wa8730 cycling genes, the ath or osa orthologues were identified and then the predicted phase was compared. While most genes are on the diagonal, which is consistent with similar phases across the species, there are some genes with different phases which could reflect that ath and osa have multiple wa8730 orthologues. For each orthogroup (OG), in each species, we computed a circular mean for the phases of the cycling genes belonging to that OG in that species. This allows us to use the OGs to map from some cycling gene in wa8730 to a ‘cycling behavior’ for orthologous genes in osa and ath. We used this mapping to create a KDE histogram plot representing the density of these pairs: phase of cycling gene, orthologous ‘cycling behavior’. This plot represents how cycling behavior in wa8730 genes related to cycling behavior in osa and ath orthologues. For example, genes that cycle and peak at dusk (11-14h) in wa8730 are associated more with dusk genes in osa and ath than with morning genes.

**Figure S13. Time of day (TOD) overrepresentation of GO terms in rice and Arabidopsis**. **A)** Arabidopsis and **B)** rice significantly overrepresented GO terms (y-axis) by TOD (x-axis). Dark color is less significant and light color is more significant.

### Supplemental Tables

**Table S1. BioNano Genomics optical map summary**.

**Table S2. *Wolffia australiana* genome size estimates by flow cytometry**.

**Table S3. Ribosomal DNA content of the Wolffia genomes**.

**Table S4. BUSCO scores across Wolffia and Spirodela genomes**.

**Table S5. Missing BUSCO gene in wa8730**.

**Table S6. Orthogroup summary (OG).**

Abbreviations: cre,*Chlamydomonas reinhardtii*; mco,*Micromonas commoda*; smo,*Selaginella moellendorffii*; atr,*Amborella trichopoda*; sly,*Solanum lycopersicum*; vvi,*Vitis vinifera*; ptr,*Populus trichocarpa*; ath,*Arabidopsis thaliana*; pab,*Picea abies*; mpo,*Marchantia polymorpha*; sp9509,*Spirodela polyrhiza*; sp7498,*Spirodela polyrhiza*; wa7733,*Wolffia australiana*; wa8730,*Wolffia australiana*; bdi,*Brachypodium distachyon*; hvu,*Hordeum vulgare*; obr,*Oryza brachyantha*; osaj,*Oryza sativa japonica*; osai,*Oryza sativa indica*; oth,*Oropetium thomaeum*; sbi,*Sorghum bicolor*; sit,*Setaria italica*; tae,*Triticum aestivum*; zma,*Zea mays*; zjn,*Zoysia japonica* ssp. nagirizaki; aco,*Ananas comosus*; egu,*Elaeis guineensis*; mac,*Musa acuminata*; ped,*Phyllostachys edulis*; peq,*Phalaenopsis equestris*; zoma,*Zostera marina*; ppa,*Physcomitrella patens*.

**Table S7. Wolffia unique orthogroup (OG) gene ontology (GO) terms**.

**Table S8. Gene in Wolffia specific orthogroups (OGs**).

**Table S9. Wolffia missing genes and the Arabidopsis annotation**.

**Table S10. Expression across all Wolffia genes**.

**Table S11. wa8730 significant cycling genes**.

**Table S12. wa8730 all elements**.

**Table S13. Significant wa8730 ELEMENTS**.

**Table S14. Wolffia TOD significant GO terms**.

**Table S15. Arabidopsis TOD significant GO terms**.

**Table S16. Rice TOD significant GO terms**.

**Table S17. TOD GO overrepresenation across Ath, Osa and wa8730**.

### Supplemental material

**Movie S1**. Tracking of the asexual propagation of *W. australiana* in culture by light microscope. Time-lapse video shows an emerging daughter frond.

