## Supplemental Figures for "Genome and time-of-day transcriptome of *Wolffia australiana* link morphological extreme minimization with un-gated plant growth"

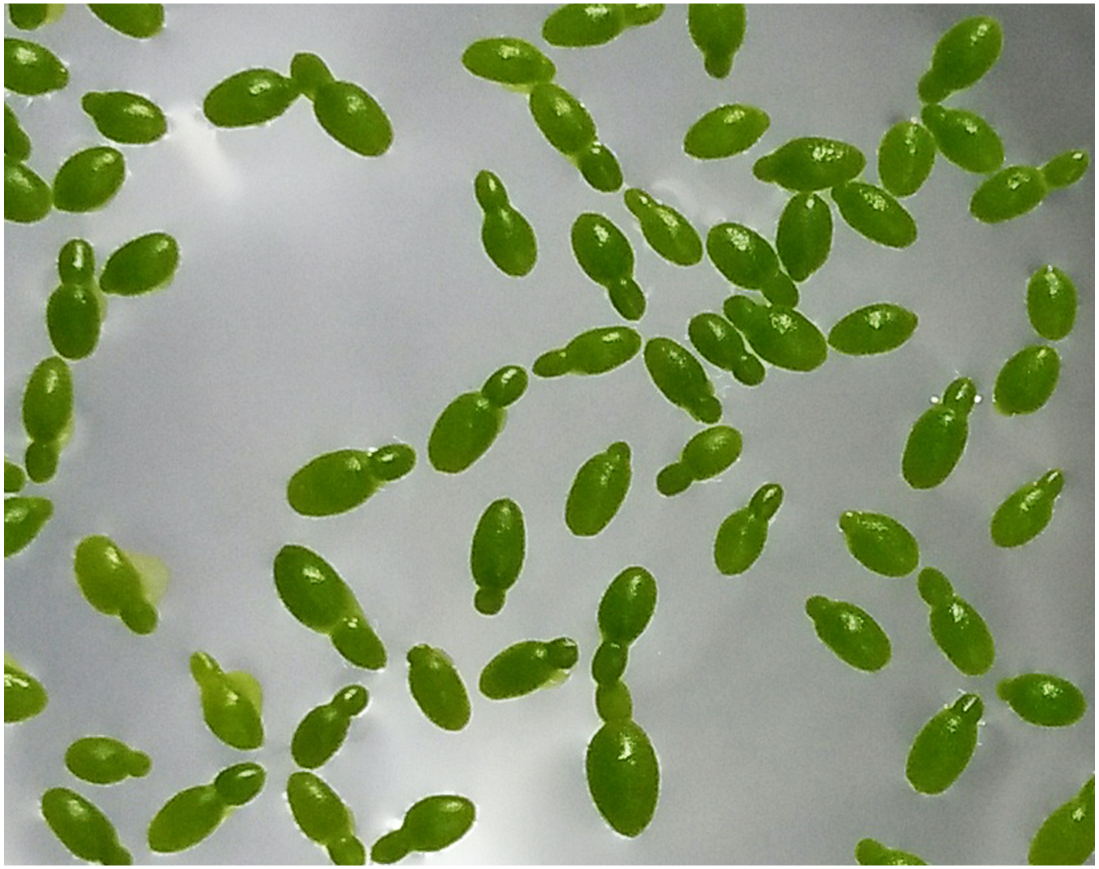

**Figure S1. Wolffia australiana under growth assay.**

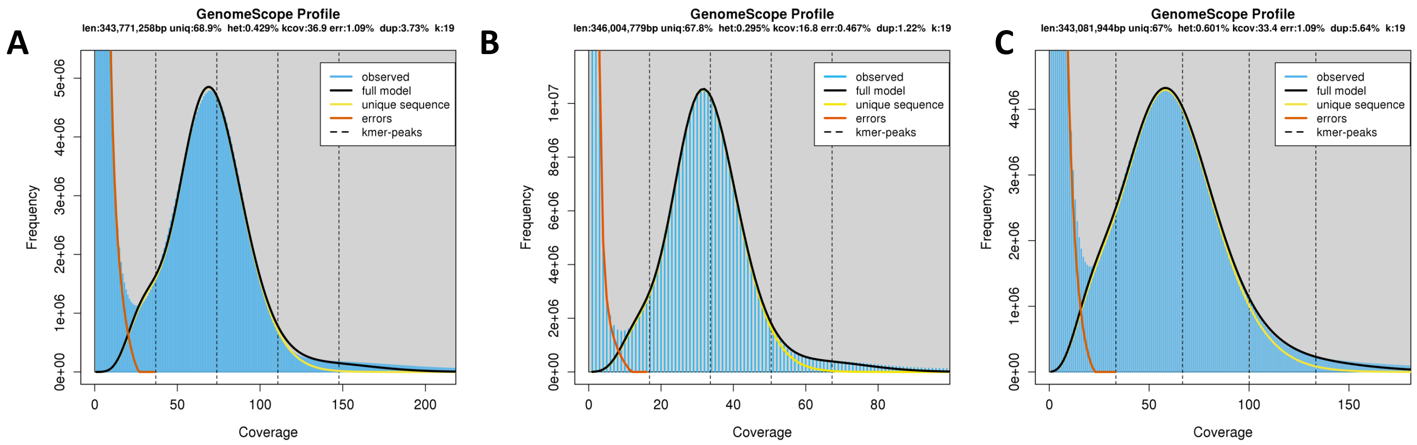


D

| | wa7733 | | wa8730 | | wa7540 | |
|---|---|---|---|---|---|---|
| property | min | max | min | max | min | max |
| Heterozygosity | 0.43% | 0.43% | 0.29% | 0.30% | 0.60% | 0.60% |
| Genome length (bp) | 343,520,172 | 343,771,258 | 345,863,147 | 346,004,779 | 342,809,947 | 343,081,944 |
| Genome repeat (bp) | 106,901,092 | 106,979,228 | 111,346,478 | 111,392,074 | 112,983,649 | 113,073,294 |
| Genome unique (bp) | 236,619,080 | 236,792,030 | 234,516,670 | 234,612,705 | 229,826,298 | 230,008,650 |
| Repeat (%) | 31.1 | 31.1 | 32.2 | 32.2 | 33.0 | 33.0 |
| Model fit | 92.21% | 96.13% | 94.73% | 98.29% | 94.74% | 98.34% |
| Read error (%) | 1.09% | 1.09% | 0.47% | 0.47% | 1.09% | 1.09% |
| Kmer | 19 | | 19 | | 19 | |

**Figure S2. Genome size estimated by Kmer (k=19) frequency.**

A

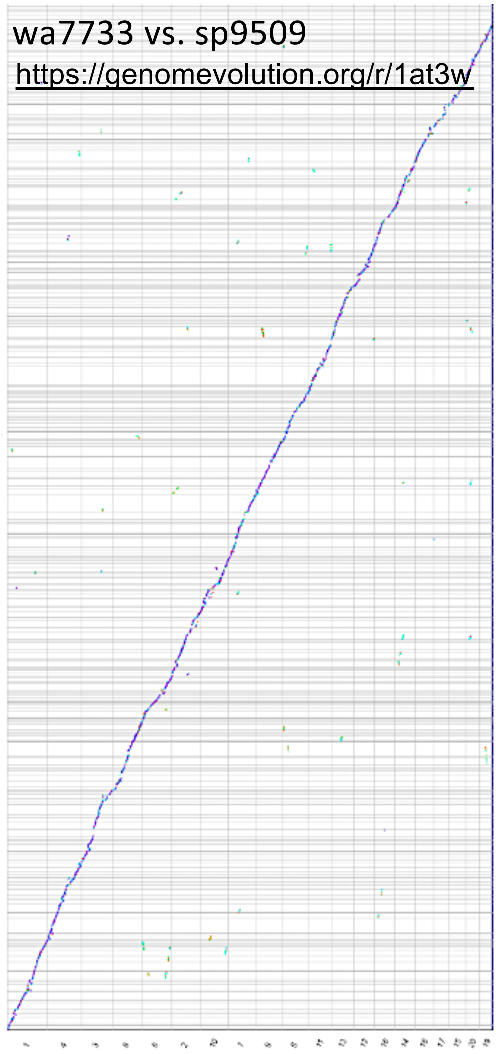


B

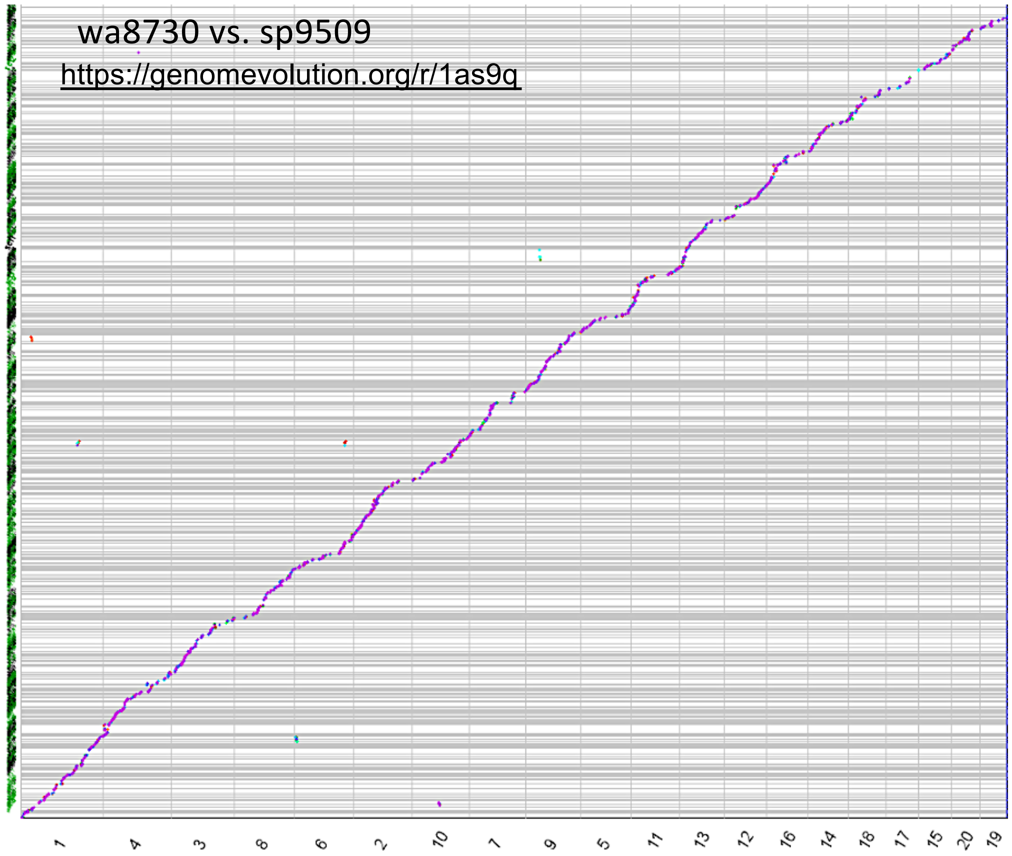


C

| | wa7733 | wa8730 |
|---|---|---|
| Total gene level syntenic blocks (>5) | 814 | 518 |
| Average number of genes per block | 10 | 7 |
| Minimum genes per block | 5 | 5 |
| Maximum genes per block | 92 | 21 |
| Total genes in syntenic blocks | 7,936 | 3,622 |

**Figure S3. Wolffia is colinear with sp9509.**

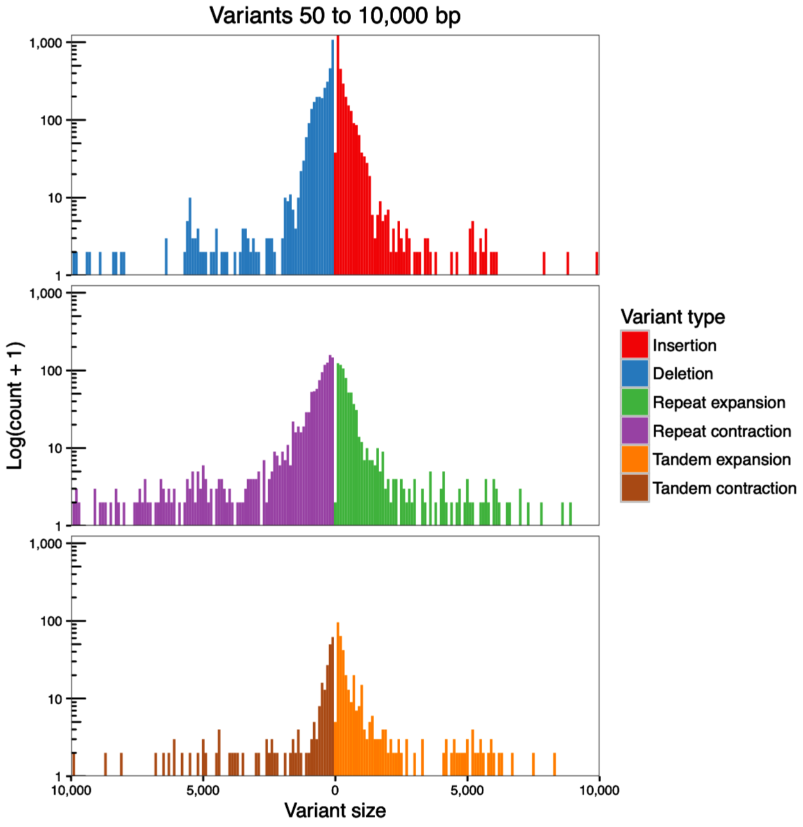


**Figure S4. Wolffia genomes differ mostly by small INDELs.**

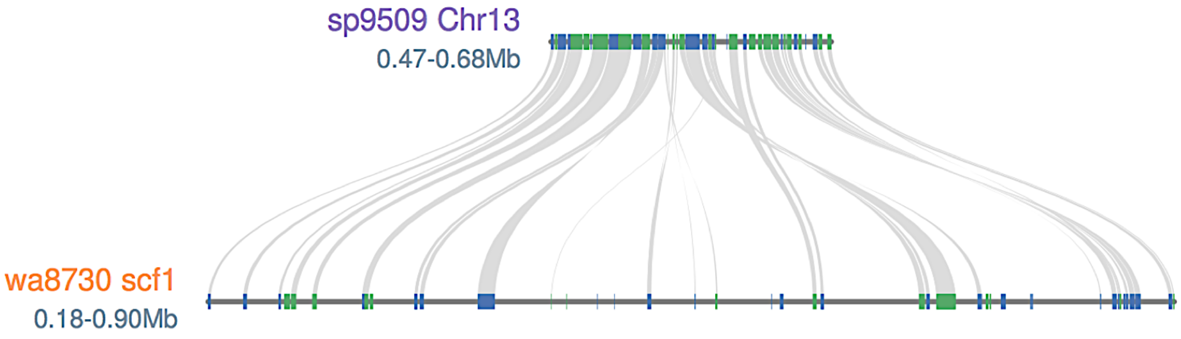


**Figure S5. Wolffia genome is colinear with sp9509.**

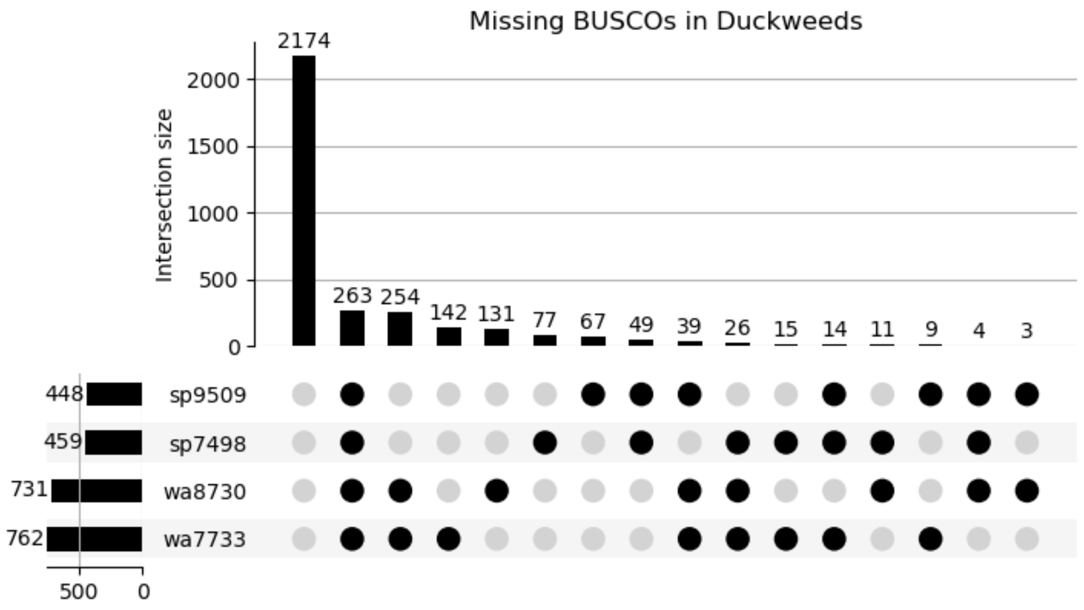


**Figure S6. Wolffia and Spirodela missing a similar set of BUSCO genes.**

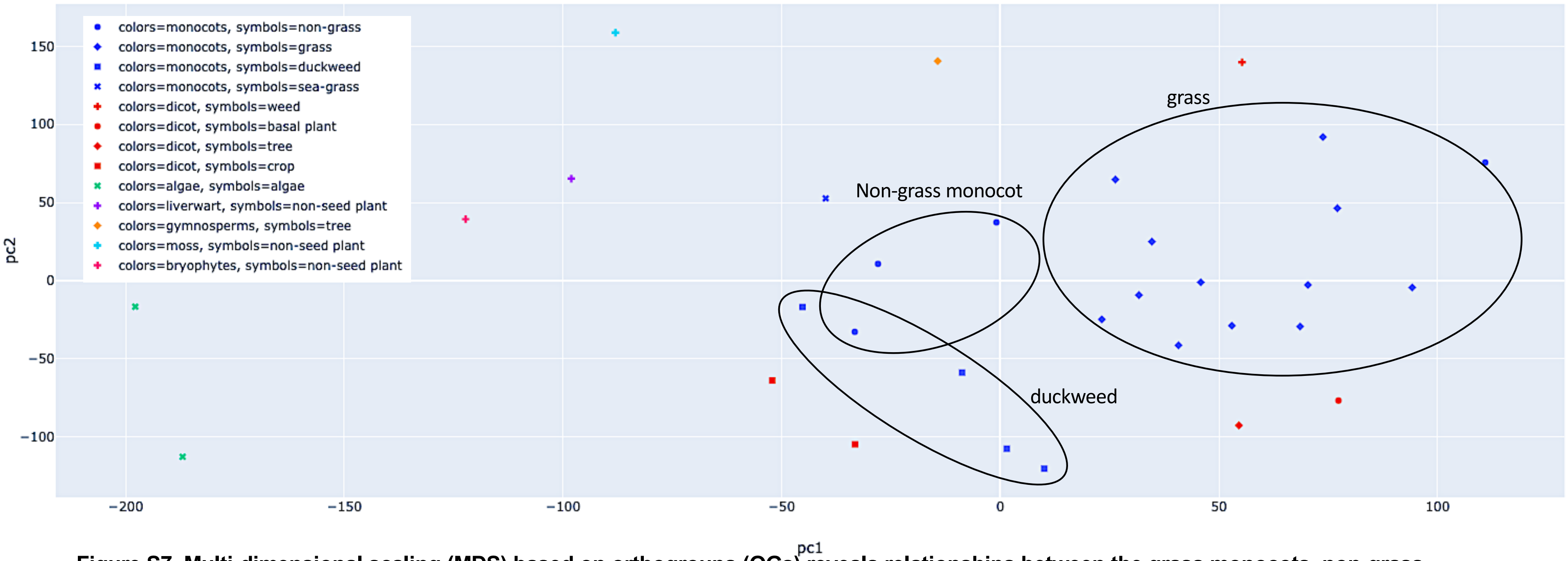


**Figure S7. Multi-dimensional scaling (MDS) based on orthogroups (OGs) reveals relationships between the grass monocots, non-grass monocots and duckweeds.**

| | Wa7733 | Wa8730 | Sp9509 | Sp7498 | zostera | arabidopsis | rice | brachy | maize |
|---|---|---|---|---|---|---|---|---|---|
| 1 | 41 | 45 | 37 | 33 | 26 | 18 | 18 | 19 | 12 |
| 2 | 20 | 20 | 19 | 18 | 17 | 14 | 12 | 13 | 13 |
| 3 | 9 | 9 | 9 | 10 | 10 | 9 | 8 | 9 | 9 |
| 4 | 6 | 5 | 6 | 6 | 7 | 7 | 5 | 6 | 8 |
| 5 | 3 | 3 | 4 | 4 | 5 | 5 | 4 | 4 | 6 |
| 6 | 3 | 3 | 4 | 4 | 4 | 4 | 3 | 3 | 5 |
| 7 | 2 | 2 | 2 | 3 | 3 | 3 | 2 | 2 | 5 |
| 8 | 2 | 2 | 2 | 2 | 2 | 3 | 2 | 2 | 3 |
| 9 | 1 | 1 | 2 | 2 | 2 | 2 | 2 | 2 | 3 |
| 10 | 1 | 1 | 1 | 1 | 2 | 2 | 2 | 2 | 2 |
| 11-15' | 3 | 3 | 4 | 4 | 6 | 7 | 6 | 6 | 7 |
| 16-20 | 2 | 2 | 2 | 2 | 2 | 4 | 3 | 4 | 5 |
| 21-50 | 3 | 2 | 3 | 4 | 4 | 8 | 8 | 8 | 9 |
| 51-100 | 1 | 1 | 1 | 1 | 1 | 2 | 3 | 3 | 2 |
| 101-150 | 1 | 1 | 0 | 0 | 1 | 1 | 2 | 1 | 1 |
| 151-200 | 0 | 0 | 1 | 1 | 0 | 0 | 0 | 0 | 1 |
| 201-500 | 0 | 0 | 0 | 0 | 0 | 0 | 0 | 0 | 1 |
| 501-1000 | 0 | 0 | 0 | 0 | 0 | 0 | 0 | 0 | 0 |
| 1001+ | 0 | 0 | 0 | 0 | 0 | 0 | 0 | 0 | 0 |

**Figure S8. Wolffia and Spirodela genes found in small gene families (orthogroups).**

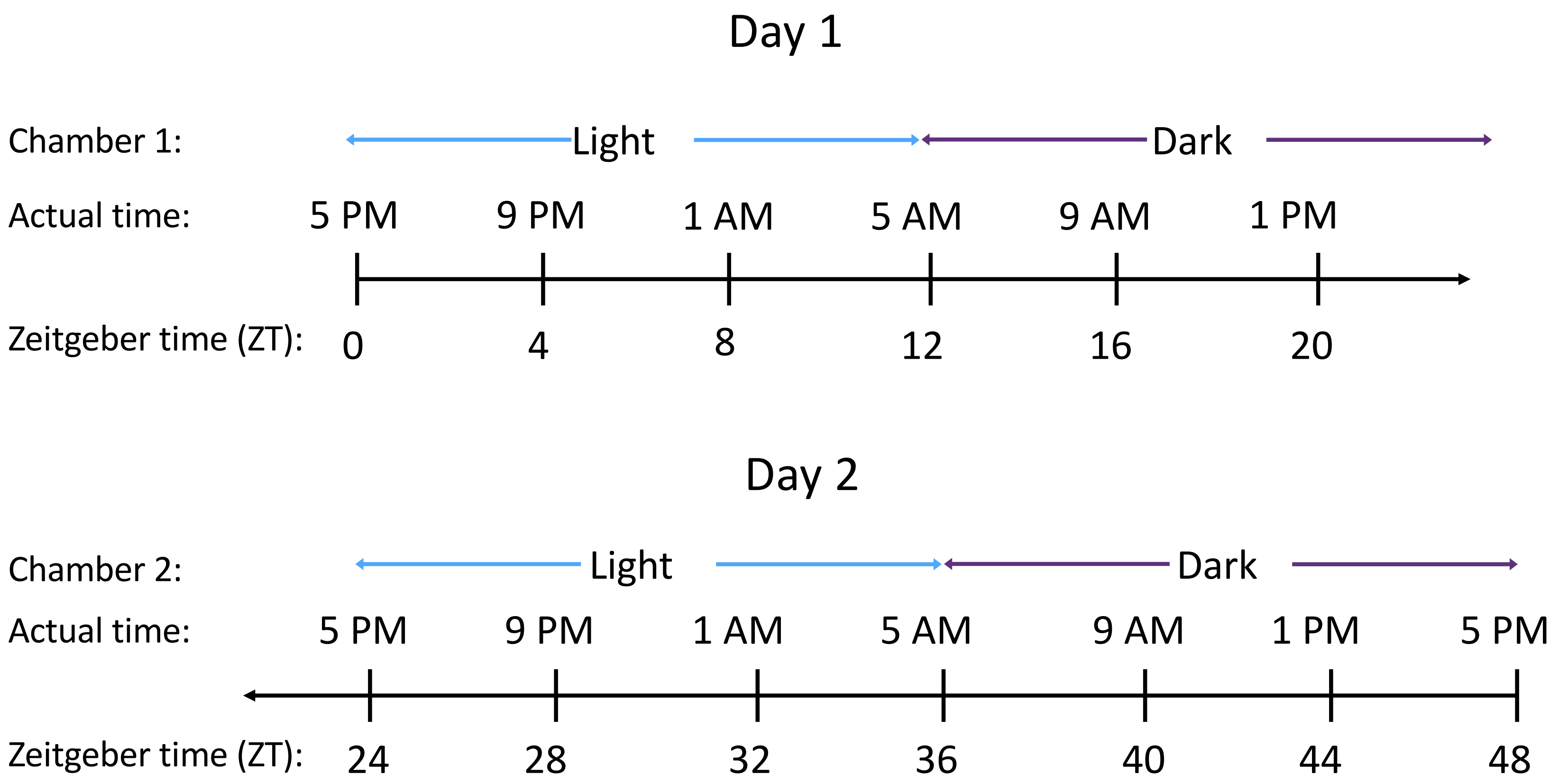


**Figure S9. Wolffia time course design.**

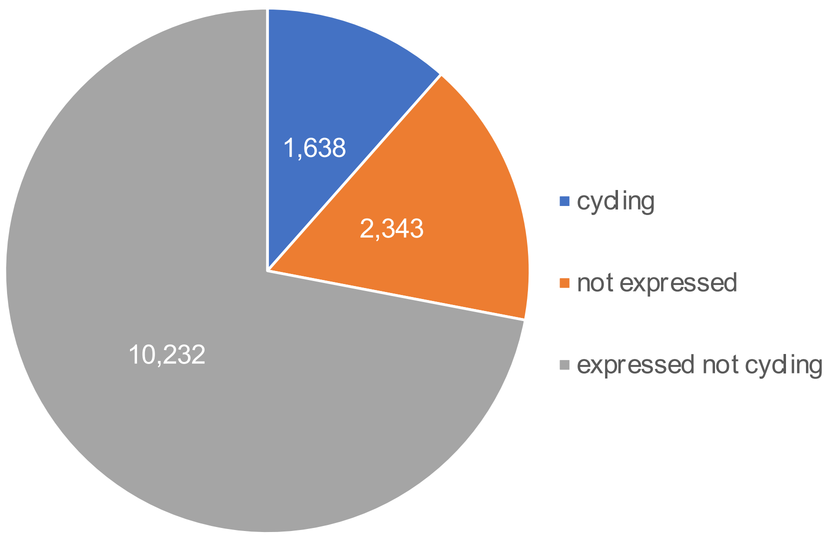


**Figure S10. Break down of expressed and cycling genes in wa8730.**

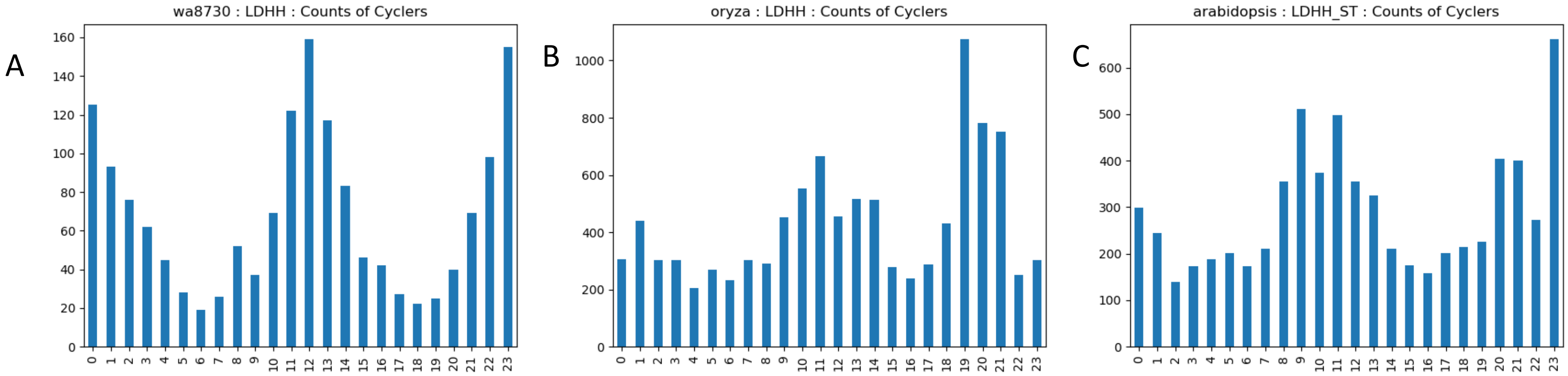


**Figure S11. Distribution of cycling genes over the day.**

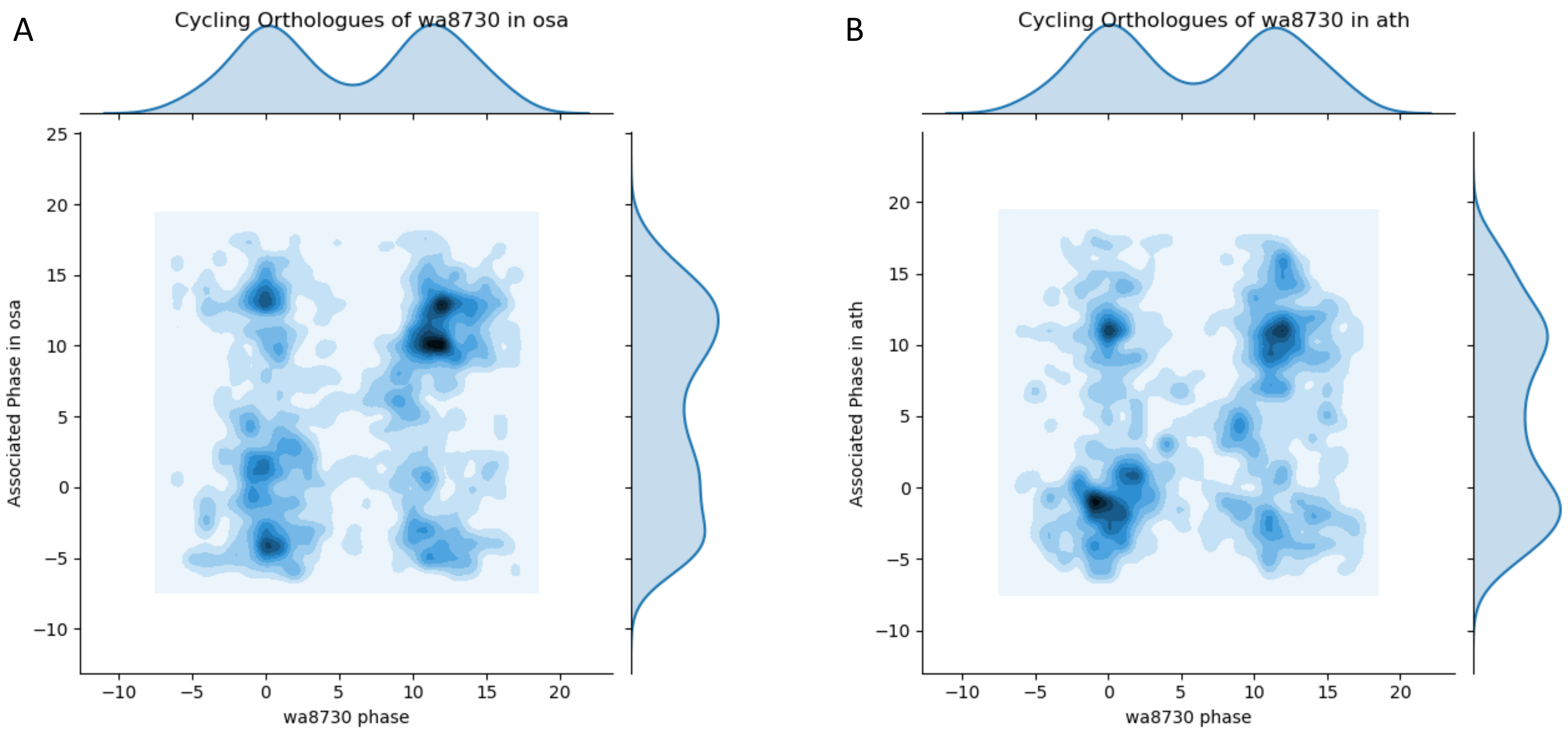


**Figure S12. Comparison of cycling genes between Wolffia (wa8730), Arabidopsis (ath) and rice (osa).**

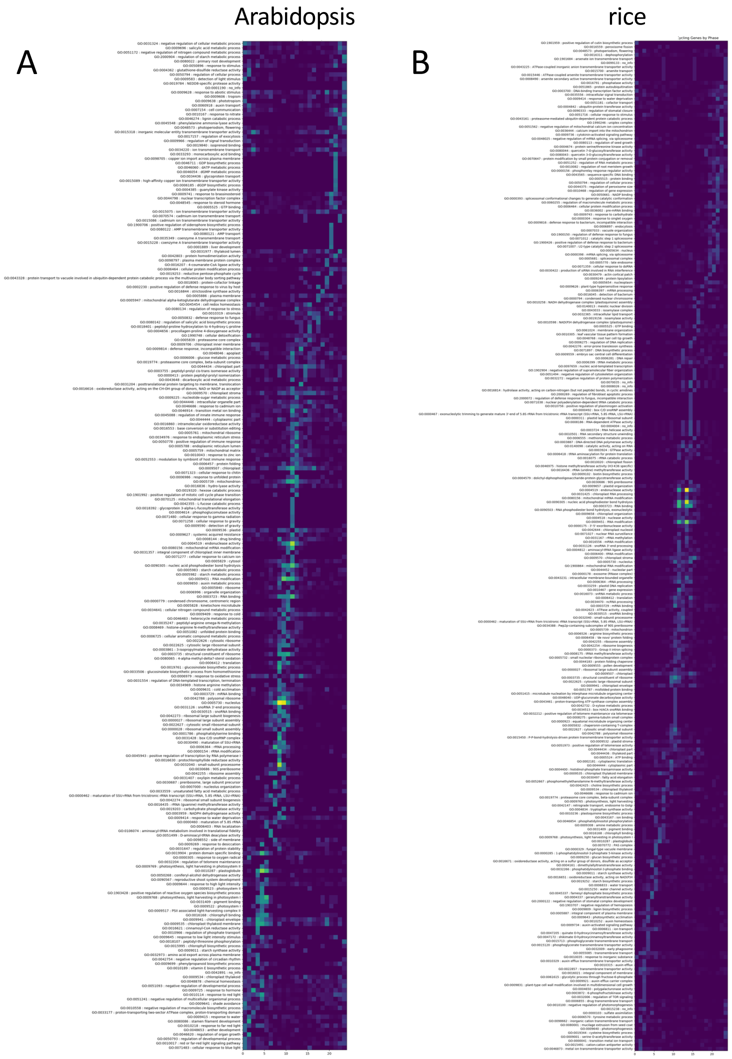


**Figure S13. Time of day (TOD) overrepresentation of GO terms in rice and Arabidopsis.**
